# Taking shortcuts: Great for travel, but not for reproducible methods sections

**DOI:** 10.1101/2022.08.08.503174

**Authors:** Kai Standvoss, Vartan Kazezian, Britta R. Lewke, Kathleen Bastian, Shambhavi Chidambaram, Subhi Arafat, Ubai Alsharif, Ana Herrera-Melendez, Anna-Delia Knipper, Bruna M. S. Seco, Nina Nitzan Soto, Orestis Rakitzis, Isa Steinecker, Philipp van Kronenberg Till, Fereshteh Zarebidaki, Tracey L. Weissgerber

**Affiliations:** Department of Psychiatry, Charité – Universitätsmedizin Berlin, Corporate Member of Freie Universität Berlin and Humboldt-Universität zu Berlin, 10117 Berlin, Germany; Bernstein Center for Computational Neuroscience, Charité-Universitätsmedizin Berlin, 10117 Berlin, Germany; Einstein Center for Neurosciences Berlin, 10117 Berlin, Germany; BIH QUEST Center for Responsible Research, Berlin Institute of Health at Charité – Universitätsmedizin – Berlin, Berlin, Germany; Berlin School of Mind and Brain, Humboldt-Universität zu Berlin, Germany; French Institute of Health and Medical Research, Interdisciplinary Institute of Social Issues IRIS (UMR 8156 CNRS - EHESS - U997 INSERM), Aubervilliers, France; Department of Life Sciences, Humboldt-Universität zu Berlin, Berlin, Germany; Berlin School of Mind and Brain, Berlin, Germany; Charité - Universitätsmedizin Berlin, Department of Neurology, Charitéplatz 1, 10117 Berlin, Germany. Charité – Universitätsmedizin Berlin; Department of Oral and Maxillofacial Surgery, Dortmund General Hospital, Dortmund, Germany; Department of Psychiatry and Psychotherapy, Campus Benjamin Franklin, Charité – Universitätsmedizin Berlin, Corporate Member of Freie Universität Berlin and Humboldt-Universität zu Berlin, Berlin, Germany; German Federal Institute for Risk Assessment (BfR), Department of Biological Safety, Berlin 10589, Germany; Former affiliation: Department of Biomolecular Systems, Max Planck Institute of Colloids and Interfaces, Potsdam, 14476, Germany. Current affiliation: Biontech SE, An D. Goldgrube 12, 55131 Mainz, Germany; Institute du Cerveau – Paris brain institute (ICM), 75013 Paris, France; École doctorale Cerveau, cognition, comportement (ED3C), Sorbonne Université Paris; Department of Psychiatry and Neurosciences, Charité – Universitätsmedizin Berlin, Corporate Member of Freie Universität Berlin and Humboldt-Universität zu Berlin, Berlin, Germany; Biological Psychology and Neuroergonomics, Berlin Institute of Technology, Berlin, 10623, Germany; Berlin Institute of Health and Neuroscience Research Center, Charité – Universitätsmedizin Berlin, Berlin, Germany; Former affiliation: Institute of Neurophysiology, Charité-Universitätsmedizin, Berlin, 10117, Germany

**Keywords:** reproducibility, transparency, protocols, methods, replication

## Abstract

Methods sections are often missing critical details needed to reproduce an experiment. Methodological shortcut citations, in which authors cite previous papers instead of describing the method in detail, may contribute to this problem. This meta-research study used three approaches to systematically examine the use of shortcut citations in neuroscience, biology and psychiatry. First, we examined papers to determine why authors use citations in the methods section and to assess how often shortcut citations were used. Common reasons for using citations in the methods section included explaining how something was done by citing a previous resource that used the method (methodological shortcut citation), giving credit or specifying what was used (who or what citation), and providing context or a justification (why citation). Next, we reviewed 15 papers to determine what can happen when readers follow shortcut citations to find methodological details. While shortcut citations can be used effectively, problems encountered included difficulty identifying or accessing the cited materials, missing or insufficient descriptions of the cited method, and chains of shortcut citations. Third, we examined journal policies. Fewer than one quarter of journals had policies describing how authors should report methods that have been described previously or asking authors to explain modifications of previously described methods. We propose that methodological shortcut citations should meet three criteria; cited resources should describe a method very similar to the authors’ method, provide enough detail to allow others to implement the method, and be open access. We outline actions that authors and journals can take to use shortcut citations responsibly, while fostering a culture of open and reproducible methods reporting.

## Introduction

Methods sections should serve several purposes, each of which requires a different level of detail (Figure 1). Well-written methods sections provide an overview of the study design and techniques used to answer the research question, help readers to evaluate the risk of bias and provide details needed to replicate the experiment. Unfortunately, methods sections are often missing critical details. The Reproducibility Project for Cancer Biology aimed to replicate high profile findings from 193 experiments in cancer research [1]. Unfortunately, none of the papers contained sufficient details to allow researchers to design and conduct a replication study [2]. When the original authors were contacted to obtain methodological details, 41% were extremely or very helpful, 9% were minimally helpful, and 32% were not helpful or did not respond [2]. An assessment of 300 fMRI studies revealed that key information, such as how the task was optimized for efficiency or the distribution of inter-trial intervals, was frequently missing [3]. Methodological details in randomized controlled trials are also frequently missing. This risk of bias for randomization sequence generation and allocation concealment was unclear, or could not be evaluated, for approximately half of the randomized controlled trials included in Cochrane reviews [4].

**Figure 1:**
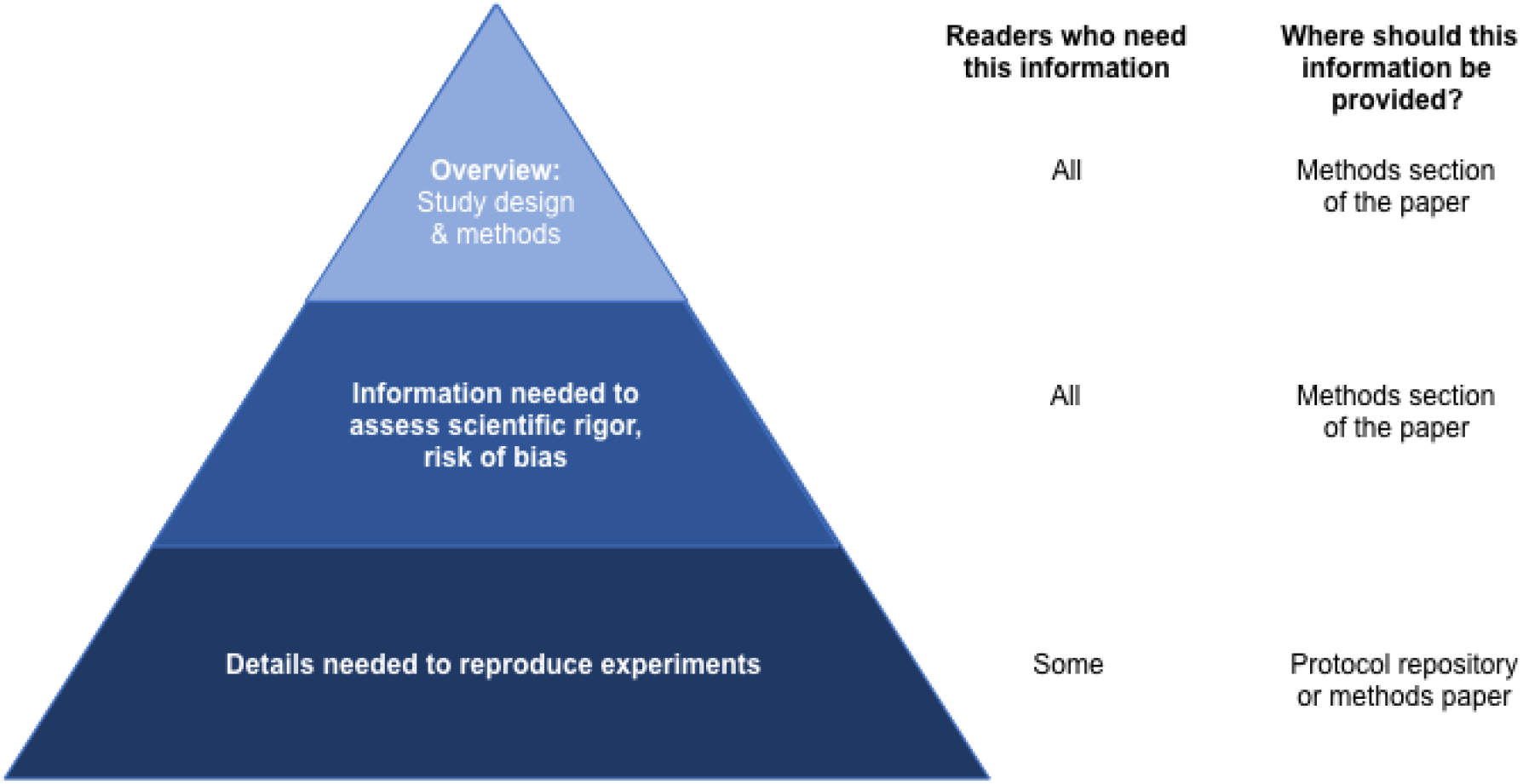
The level of methodological detail required depends on the reader. All readers need an overview of the study design, methods used to answer the research question, and information needed to assess scientific rigor and the risk of bias. These details should always be presented in the methods section of the paper. While fewer readers need details required to reproduce the method, these readers are particularly important because they are most likely to perform follow-up experiments. Sharing this level of detail in the methods section is often difficult; therefore, this information should be shared in a protocol repository or methods paper.

A methodological shortcut citation is a citation that is used to replace a full description of a method or a part of a method, under the assumption that the resource being cited fully describes the method. Shortcut citations are sometimes accompanied by phrases like “as described previously” or “see (citation) for details”. Methodological shortcut citations are one factor that could affect readers’ ability to reproduce an experiment.

While some scientists and editors view the use of shortcut citations as a good practice [5, 6], others worry that these shortcuts may adversely affect reproducibility [7]. Ideally, shortcut citations should reference state of the art resources that describe the method in detail. This may include protocol papers, diagnostic guidelines, or original research articles with detailed methods. In practice, anecdotal reports show that shortcut citations can cause problems [8, 9]. Readers may be unable to access the cited paper. The cited paper may not contain methodological details or may itself use a shortcut citation instead of describing the methods. Those in favor of shortcut citations note that authors don’t waste time repeating details that have been written elsewhere and avoid potential copyright issues that might emerge if one copied methods from another publication. Those who are skeptical of shortcuts argue that the cited paper may no longer be state of the art or may not accurately describe the procedures of the authors who cite it. Furthermore, consulting cited resources to obtain details needed to interpret the study is time consuming. There are also disagreements about how shortcut citations should be used. While some authors cite the paper that introduced the method to give its creators credit, others cite the paper whose methods most closely resemble their own. Some journals require authors to use shortcut citations to avoid repeating published methods, or incentivize this practice through strict word limits, whereas other journals have excluded the methods section from word limits to encourage detailed reporting [10].

Our meta-research study used three approaches to systematically examine the use of methodological shortcut citations in neuroscience, biology and psychiatry. First, we examined papers to determine why authors use citations in the methods section and to assess how often shortcut citations were used. Next, we reviewed shortcut citations for 15 papers to determine what can happen when readers follow these citations to find methodological details. Third, we reviewed journal policies related to shortcut citations and methodological reporting.

## Methods

This study was performed as part of a participant guided, learn-by-doing course [11], in which graduate students in different fields at four Berlin universities learned meta-research skills by working together to design, conduct and publish a meta-research study.

We conducted three distinct but related studies.

1. **Prevalence Study:** This study examined the reasons why authors use citations in the methods section of papers and assessed how often shortcut citations were used. Additional data collected included the number of citations per shortcut and the years in which papers cited as shortcuts were published.
2. **Case Series:** This study examined problems that may occur when readers consult shortcut citations to find further details of the study methods.
3. **Journal Policy Study:** This study examined journal policies related to shortcut citations and methodological reporting.

Each study examined three fields; biology, psychiatry and neuroscience, to improve generalizability. These fields were selected based on the study teams’ expertise. The abstraction protocols, data and code for the prevalence study and journal policy studies were deposited on the Open Science Framework (RRID:SCR_003238; https://osf.io/d2sa3/?view_only=e130a3a44a9d4b148b8ea3ed61646e7e [12]).

## Prevalence Study

### Systematic Review

We followed all relevant items in the PRISMA guidelines [13]. Items that only applied to meta-analyses or were not relevant to literature surveys were not followed. Ethical approval was not required.

### Journal screening

Our sampling frame included top journals that publish original research in each field. Journals for each category were ranked according to 2019 impact factors listed for the specified categories in Journal Citation Reports. We excluded journals that did not publish original research or did not publish a March 2020 issue. Fewer journals were used for biology (n = 15), compared to neuroscience and psychiatry (n = 20), due to the large number of publications in some biology journals.

### Search strategy

Articles were identified through a PubMed search. We performed a supplemental Web of Science search to identify articles published in journals that were not indexed in PubMed. The full search strategy is available on the OSF repository [12].

### Inclusion and exclusion criteria

The study included all full length, original research articles that included a methods section published in each included journal between March 1 and March 31, 2020. Among journals that publish print issues, we examined all articles included in print issues of the journal that were published in March 2020 (Table S1, Table S2, Table S3, Figure S1). Articles for online journals that did not publish print issues were included if the publication date was between March 1 and March 31, 2020. Articles were excluded if they were not full-length original research articles, did not contain methods sections, or if an accepted version of the paper was posted as an “in press” or “early release” publication; however, the final version did not appear in the March issue.

### Screening

Screening for each article was performed by two independent reviewers (Biology: PvK, SC, ADK, VK; Neuroscience: KS, FZB, SA; Psychiatry: IS, OR, UA) using Rayyan software (RRID:SCR_017584), and disagreements were resolved by consensus discussions between the two reviewers. A list of articles was uploaded into Rayyan. Reviewers independently examined each article and marked whether the article was included or excluded.

### Abstraction

All abstractors completed a training set of 35 articles before abstracting data. Data abstraction for each article was performed by two independent reviewers (Biology: SC, ADK, VK, PvK; Neuroscience: KS, FZB, SA; Psychiatry: BL, OR, ALH). When disagreements could not be resolved by consensus, ratings were assigned after a group review of the paper. Eligible manuscripts were reviewed in detail to evaluate the following questions according to a predefined protocol (available at: https://osf.io/d2sa3/?view_only=e130a3a44a9d4b148b8ea3ed61646e7e [12]).

The following data were abstracted:

1. Is the paper related to the SARS-CoV-2 pandemic?
2. Does the paper include additional methodological details in the supplement?
3. Does the paper reference a repository or repositories as a shortcut for *any* method used? If so, what is the name of the repository? (Note that code used to analyze data was considered a method).

Abstractors then reviewed each citation in the methods section, including citations listed in STAR (Structure Transparent Accessible Reporting) methods tables [14]. Some journals use these tables to provide an overview of all of the key reagents used in the study. Notations or hyperlinks that appeared only in the text, and not as entries in the reference list (e.g. company names), were not counted as citations. Citations in the supplemental methods were not evaluated. Multiple papers cited as part of the same reference were treated as a single citation (e.g. two citations appearing within the same set of brackets). Papers cited in different locations in the same sentence were treated as different citations.

Each methodological citation was classified into one of the categories outlined in Table 1, according to the inferred purpose of the citation. The “How (Explain a method)” and “Other” categories could be shortcut citations, whereas other categories could not. Each citation in these two categories was classified as a probable shortcut, possible shortcut, or no shortcut. The conceptual goal was to distinguish between situations where readers would likely need to consult the cited paper to implement the method (probable shortcut), situations where the reader may need to consult the shortcut citation to implement the method (possible shortcut), and cases where the reader would not need to consult the shortcut citation to implement the method. However, these conceptual definitions are highly subjective and depend on the reader’s knowledge of the reported methods. We therefore used the syntactic definitions described below to classify shortcut citations.

∘ **Probable shortcut:** The sentence that includes the shortcut citation is the only description of the method. Additional details are not provided in the following sentences or elsewhere in the methods section.
∘ **Possible shortcut:** Additional details of the cited method are provided in the sentences following the sentence containing the shortcut citation, or elsewhere in the methods section.
∘ **Not a shortcut:** A reader would not need to consult the cited paper to implement the method. This rare category was generally used when the citation referred to concentrations, parameters or other details that were fully specified in the sentence where the shortcut citation appeared.

**Table 1:** Reasons for citations in the methods section

| Category | Example | Potential shortcut |
| --- | --- | --- |
| **How (Explain a method):** The citation was intended to explain how something was done. This category included four subcategories for specific types of citations (1. General, 2. Protocol, 3. Prior publication describing the study design and/or results, and 4. Guideline or manual). | Liver samples were collected, sectioned and frozen as described previously (citation). | yes |
| **Who or what (Give credit or specify what was used):** The citation is used to give credit to the group that created the method, tool, resource or substance, or to specify exactly what method, tool, resource or substance was used. The citation is not used to explain how the method was performed, or how the tool, resource or substance was used. This category includes three sub-categories (1. Software, 2. Atlas, 3. Other). | Analyses were conducted using R (citation of R). | no |
| **From where (Source of materials):** The citation is used to show where a substance or organism was obtained from, not how the substance or organism was created. | Mice were obtained from the Smith lab (citation). | no |
| **Why (Provide context or a justification):** The citation refers to previous studies to provide context. This includes citations that compare the authors' results with results from previous studies, demonstrate that others have used a similar approach, or explain why the authors chose a particular method, model, formula, etc. The citation is not intended to provide insight into how the existing study was conducted. | In accordance with previous studies (citation), we observed that the optimal treatment dose was 20 ml. | no |
| **Formula or value:** The citation referred to a formula or specified a value for a parameter that was used. The formula or value had to be stated in the article, so that the reader would not need to look up this information in the cited resource. The citation was not used to explain how that parameter was calculated, how the formula was derived, or why the parameter or formula was chosen. | Drug cost \$X for drug Y was obtained from publicly available reports (citation). | no |
| **Other:** Citations that do not fit into the other categories. |  | yes |

A complete protocol with examples is available in the online repository.

Two independent reviewers (SA, UA) abstracted the following data for each paper:

- The minimum and maximum number of citations within each probable shortcut
- The minimum and maximum number of citations within each possible shortcut
- The publication year of the youngest and oldest probable shortcut citations
- The publication year of the youngest and oldest possible shortcut citations

## Case Series

### Selection of articles

In each field, papers from the prevalence study were divided into quintiles based on the total number of shortcuts (possible + probable). Ten articles from each quintile were randomly selected using a computer algorithm and placed on an ordered list. Potential reviewers for the case series identified articles on the list that fell within their area of expertise. We then chose the first article in each list that had a self-identified expert evaluator. This approach was used to select 15 parent articles (one article per quintile per field).

### Data abstraction

Each reviewer carefully examined all papers or materials cited as shortcuts to determine whether they could locate the cited content. While articles were almost always accessible, reviewers were sometimes unable to access other types of cited resources. Books were only checked if they could be accessed online, as students could not visit libraries due to COVID-19 pandemic restrictions. Newer versions of books and manuals were reviewed if older, cited versions were unavailable or inaccessible.

For each cited article or resource that was found, the reviewer documented the cited material type (paper, protocol, book, etc.), publication year, whether the article or resource was open access or behind a paywall, and any other problems encountered while searching for information on the cited method. Additionally, reviewers noted whether book citations included chapters or pages. Finally, reviewers also noted whether the article or resource contained an adequate description of the method cited in the parent article. If the methodological description was not adequate and the shortcut citation also used a shortcut citation to explain the method cited in the parent paper, then the reviewer repeated the abstraction process for each new shortcut citation, adding these new shortcuts as additional steps in the shortcut citation chain. Abstraction was complete when the reviewer either found a complete and comprehensive description of the method or reached a dead-end in the chain of shortcut citations. Dead ends included an inability to locate the cited article or resource, or an article or resource that did not describe the cited method.

A second reviewer assessed the accuracy of all abstracted information. A graphic illustrating the number of shortcuts and chains of shortcut citations was created for each parent paper.

## Journal policy study

We examined policies of all eligible journals listed in the Journal Citation Reports 2019 ranking for neuroscience, psychiatry and biology.

### Inclusion and exclusion criteria

Journals that publish original research were included, whereas journals that only publish review articles, book series, correspondence, perspectives, or editorials were excluded. Methods journals were excluded, as these journals often require authors to report extensive methodological details. Journals were also excluded if they did not have a website, had suspended publishing activities, or planned to cease publishing in 2021.

### Search strategy

Journal webpages were examined to confirm that journal policies were accessible. Electronic searches were performed using the terms “[journal name]”, “journal citation reports ranking”, “author guidelines”, “journal policy”, and “impact factor”. When journals with similar names were identified, impact factors were used to confirm that the correct journal was selected.

### Screening

Each journal was screened by two of the three independent reviewers (BS, KB, NS) to determine whether the journal was eligible according to the pre-specified criteria listed above. Discrepancies were resolved by consensus. Information on whether the journal publishes original research articles was assessed through the journal descriptions (e.g. “About the Journal” or “Aims and Scope”), or by determining whether the submission guidelines listed original research as an article type. If this information was inconclusive, the two most recent issues were examined to determine whether the journal published original research.

### Abstraction

Training was performed on the top 20 eligible journals for each category prior to data abstraction. Data were collected by two of the three independent abstractors (BS, KB, NS). When disagreements could not be resolved by consensus, discrepancies were resolved through discussion with the third reviewer. An additional study team member (TLW) was consulted to resolve complex discrepancies. A trained abstractor abstracted data from non-English journal webpages with the help of a native speaker.

Webpages containing author and or submission guidelines were identified by looking for the following terms in the website menu, or by using the website search function: “policy”, “policies”, “author”, “author/s guidelines”, “author/s instructions”, “submission guidelines”, “submit your article”, “recommendation”, “about”, “publish”. Each webpage section was visually examined using the search terms “method”, “methods”, “experiment”, “reproducibility”, “replicate”, “replication”, “repository”, “repositories”, “self-plagiarism”, “supplementary”, “supplementaries”, “citation”, “protocol”, “journal protocol”, “scientific society”, and “society”.

Reviewers determined whether the instructions for authors or journal policies addressed the following points:

1. Explicitly asked authors to provide sufficient methodological details to allow others to reproduce the experiment.
2. Specified how authors should describe methods that have been reported elsewhere (e.g. provide a citation instead of describing the method, briefly summarize the method and provide a citation, etc.).
3. Encouraged authors to use supplemental files, protocol repositories or protocol journals to explain their methods.

Reviewers assessed whether this information was found in the material and methods section of the author guidelines or journal policies, in other sections of the author guidelines, or elsewhere on the journal’s website.

Reviewers also consulted the journal website to determine whether the journal was affiliated with a scientific society, and whether the journal endorsed the TOP (Transparency and Openness Promotion) guidelines. The Center for Open Science list of journals that have implemented the TOP guidelines (https://www.cos.io/initiatives/top-guidelines, https://osf.io/mwxb3/) was consulted to identify journals that endorsed TOP, but did not provide this information on their website.

### Data analysis

Code for data figures and color schemes was adapted from a previous paper [15]. Summary statistics were calculated using Python (RRID:SCR_008394, version 3.7.7, libraries NumPy 1.18.5 [16], Pandas 1.2.4 [17] and Matplotlib 3.4.1 [18, 19]). Charts were prepared with a Python-based Jupyter Notebook (Jupyter-client, RRID:SCR_018413 [20], Python version 3.7.7, libraries NumPy 1.18.5 [16], Pandas 1.2.4 [17] and Matplotlib 3.4.1 [18, 19]) and assembled into figures with vector graphic software.

## Results

### Prevalence Study

#### Study sample

The study sample consisted of 224 articles with 2,754 methodological citations in neuroscience, 431 papers with 6,227 methodological citations in biology and 160 papers with 1,869 methodological citations in psychiatry (Figure 2a, Figure S1). The sample contained few publications related to COVID-19 (neuroscience: n = 0, 0%, biology: n = 2, 0.5%, psychiatry: n = 0, 0%).

**Figure 2:**
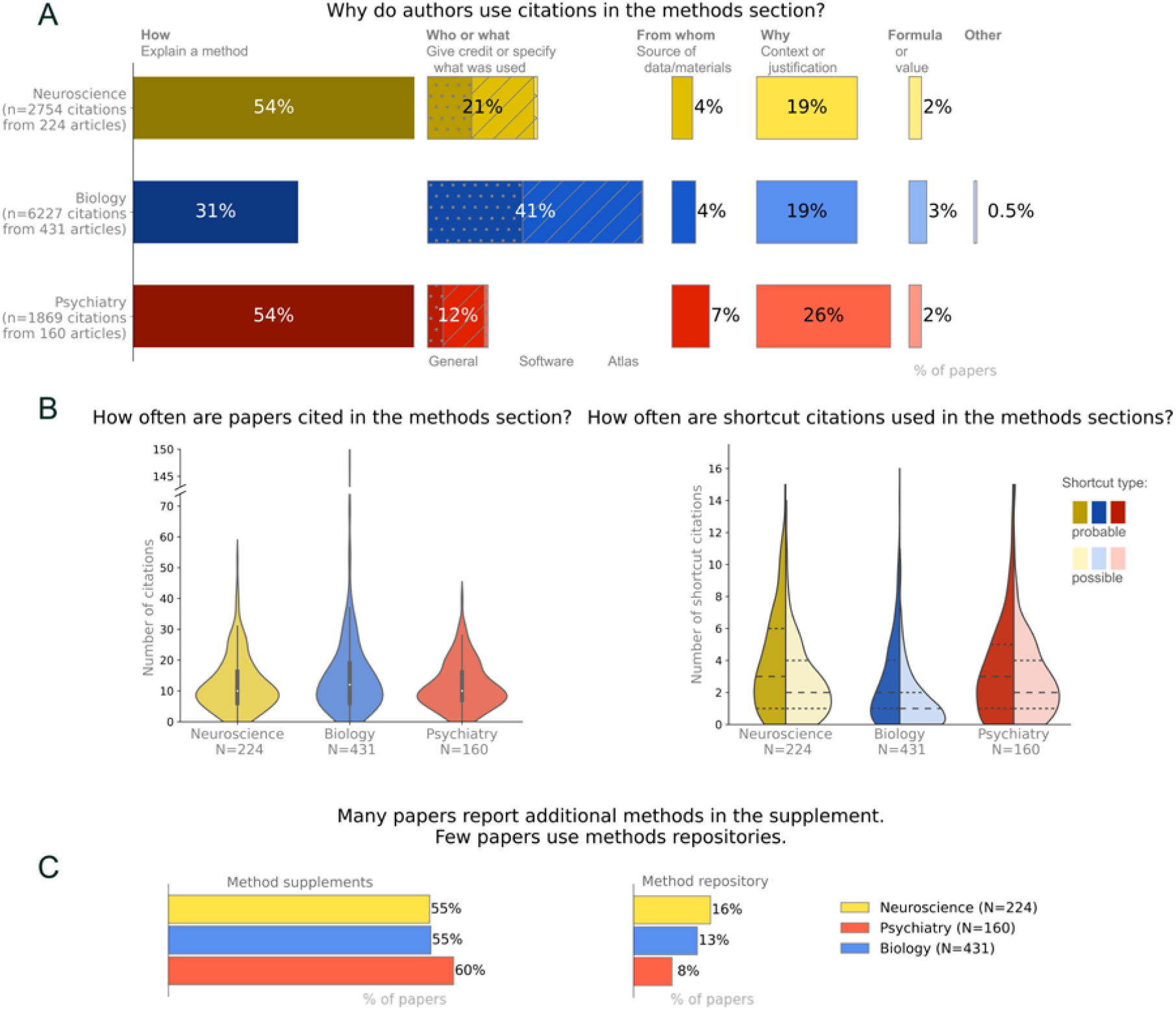
Understanding the use of citations in the methods sections, methods supplements and methods repositories. (A) The most common reasons why authors cite papers in the methods section are to explain how a method was performed, give credit or specify what was used (who or what), or provide context or a justification (why). These numbers should be regarded as approximations. Small changes in the wording or position of the reference could alter the categorization, as could variations in reader expertise (see limitations). (B) Citations and shortcut citations often appear in the methods section of published papers. (C) Methods are often shared in supplements but are less likely to be deposited on methods repositories. Percentages may not total 100% due to rounding errors.

#### Use of citations in the methods section

Citations were common in the methods sections of published papers (Figure 2b, left panel). The median number of citations in the methods section was 10 [interquartile range: 6, 16.25] in neuroscience, 12 [6, 19] in biology, and 10 [7, 16] in psychiatry.

In neuroscience and psychiatry, 54% of citations in the methods section were used to explain study methods (How citations, Figure 2a). Citations were also commonly used to give credit or specify what was used (Who or what, 12-21% of papers), and provide context or a justification (Why, 19-26%). In biology, the most common reason for citing a paper in the methods section was to give credit or specify what was used (Who or what, 41%), followed by explaining a method (How, 31%) and providing context or a justification (Why, 19%). Citations specifying the source of data or materials (From where), or referring to a formula or value, were uncommon in all three fields. Citations that did not fit into any of these categories were rare.

Depending on the field, 55 and 60% of papers provided some methodological information in supplemental files whereas only 8-16% provided methodological information in a repository (Figure 2c). While reviewers did not systematically collect data on the content or quality of information in the supplemental methods, reviewers’ subjective impression was that supplemental methods were rarely detailed. Common examples of supplemental methods included tables that list primers or sequences or provided basic information on study participants. Methodological information in repositories included a mixture of study protocols and code for data analysis. The most common repository was GitHub, followed by ClinicalTrials.gov, OSF and FigShare (Table S4). Other repositories were rarely used in this dataset.

#### Shortcut citations

Methodological shortcut citations were common in all three fields, with 96% of neuroscience papers, 90% of biology papers and 92% of psychiatry papers containing at least one possible or probable shortcut. Figure 2b (right panel) shows the median number of possible and probable shortcuts for each field. Summary statistics are reported in Table S5.

Reviewers assessed the age of the newest and oldest methodological shortcut citation in each paper. Figure 3a shows the number of probable (left side of violin plot) and possible (right side of violin plot) shortcut citations in each field. The median age of the newest shortcuts citation ranged between 3 and 5 years, (Figure 3a, left panel), whereas the median age for the oldest shortcut citations ranged between 10 and 24 years (Figure 3a, right panel). Summary statistics for Figure 3a are reported in Table S6.

**Figure 3:**
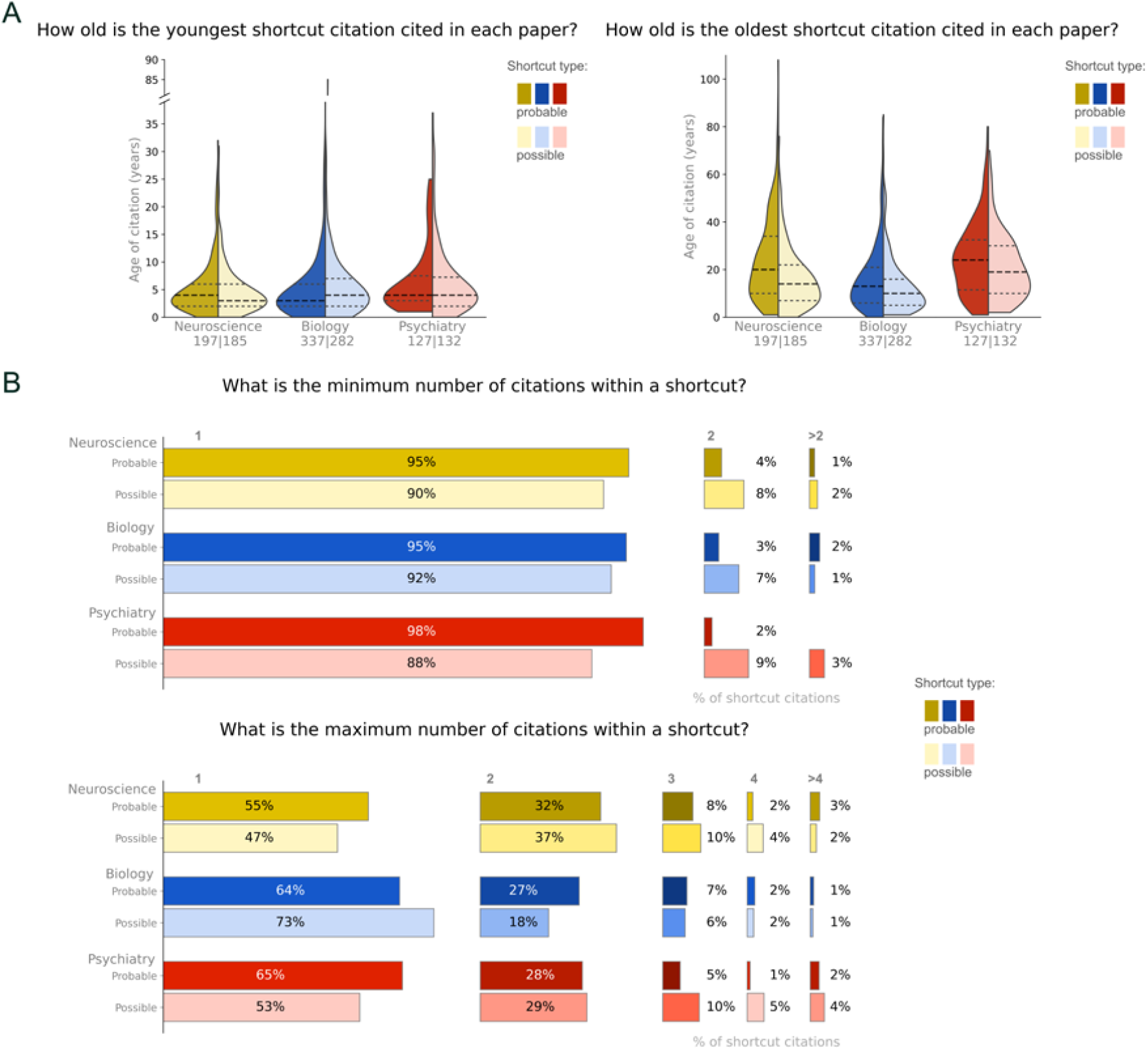
Age and number of resources cited within a shortcut citations. (A) The youngest shortcut citation cited in a paper is typically less than 10 years old. The age of the oldest shortcut citation is much more variable, and many of these citations were published more than a decade ago. The line with long dashes represents the median, whereas the lines with short dashes represent the 25^th^ and 75^th^ percentiles. The number of papers appears below the field name (n for probable shortcut citations/n for possible shortcut citations). (B) Shortcut citations typically cite one or two resources. Percentages may not total 100% due to rounding errors.

The minimum number of resources cited within a shortcut was one for 88 to 98% of papers (Figure 3b, upper panel). The maximum number of resources cited within a shortcut was typically one or two (Figure 3b, lower panel).

## Case Series

Supplemental figure S2 illustrates the process of finding cited methodological information for each article. In many cases, reviewers were able to locate additional information. However, the case series highlighted five types of issues that arise when readers follow shortcut citations in search of detailed methods (summarized in Figure 4). The first problem was an inability to identify the cited material due to incomplete or inaccurate citations (e.g. incorrect author names, years or DOIs) or dead website links. In some cases, reviewers could not find evidence that the cited source existed. The second problem was accessing the cited source. PDFs for some older articles were difficult or impossible to obtain. Many articles, supplemental files and books were not open access. Subscriptions vary among institutions, and even scientists from well-funded institutions may have difficulty accessing paywalled articles. Recent Elsevier publications, for example, were inaccessible to study team members from Berlin institutions due to ongoing contract disputes between German universities and Elsevier. The third problem was that the cited method was difficult or impossible to find. Some textbook citations failed to provide specific chapters or pages, making it difficult to locate the cited content. In one case, the cited source was in a different language. The fourth problem was that the cited method was not adequately described. Examples included cited sources that didn’t describe the method, provided the same details as the citing paper or provided a description that was no longer state-of-the-art. The fifth problem was that in many cases, reviewers had to follow a chain of shortcut citations to locate a complete methodological description. Following a citation chain is time-consuming and readers may have difficulty identifying and accessing the cited material at each step in the chain.

**Figure 4:**
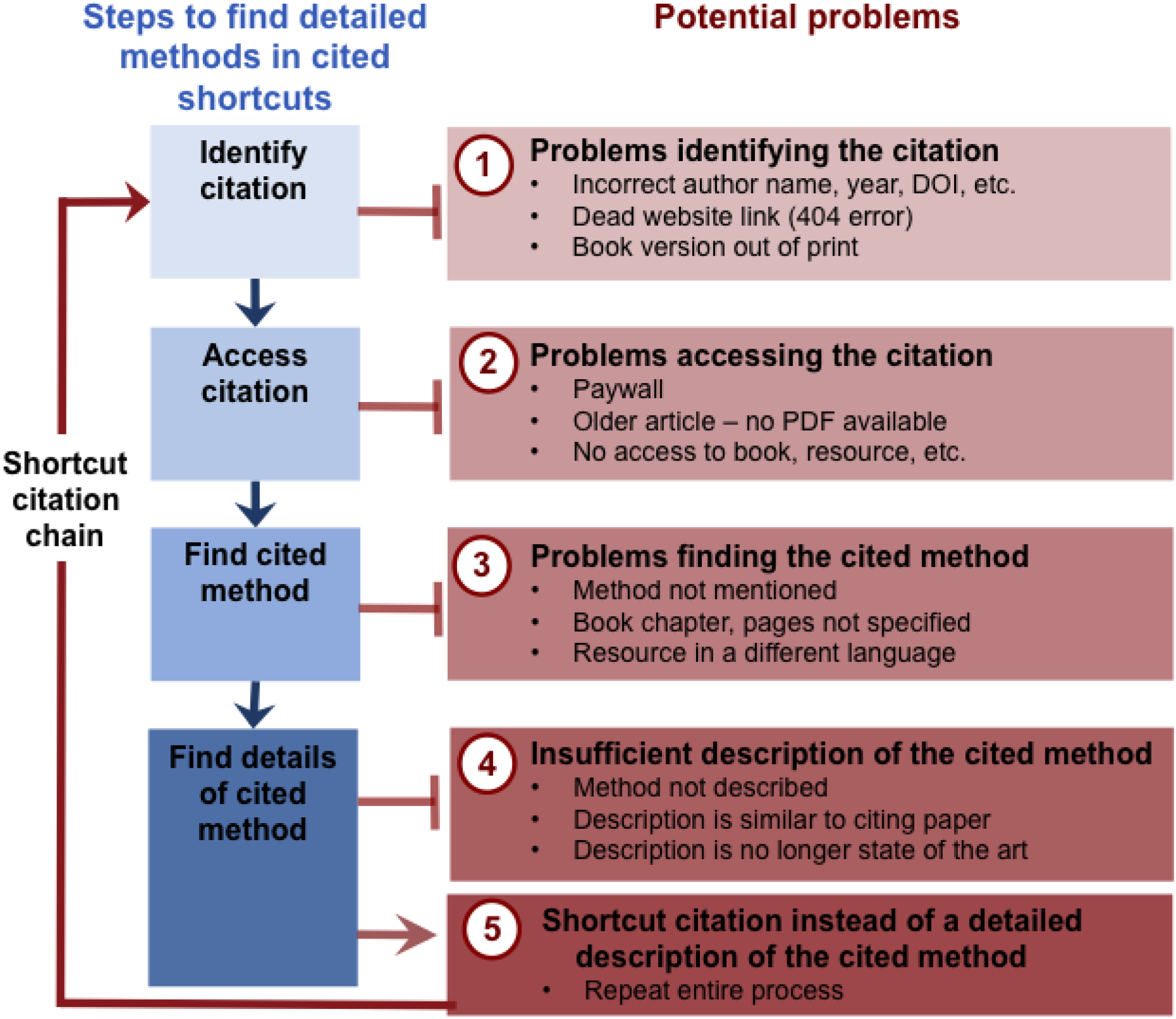
Problems that arose when searching shortcut citations for detailed methods. While methodological shortcut citations can be used effectively, reviewers encountered some problems when consulting shortcut citations to find details of cited methods. These included problems identifying the citation, problems accessing the citation, problems finding the cited method within the shortcut citation, and an insufficient description of the cited method. Chains of shortcut citations, in which the cited shortcut citation also used a shortcut citation to describe the cited method, were common.

## Journal Policy

### Study sample

Among the 519 journals screened, 465 were eligible for the study (Figure S3; 244 neuroscience journals; 76 biology journals; 145 psychiatry journals). Twenty-one journals appeared on the neuroscience and psychiatry lists and were included in both groups.

### Policies on details needed to reproduce experiments

Policies that explicitly instruct authors to provide sufficient methodological details to allow others to reproduce the experiment were found in 40% of neuroscience journals, 18% of psychiatry journals and 44% of biology journals (Figure 5a).

**Figure 5:**
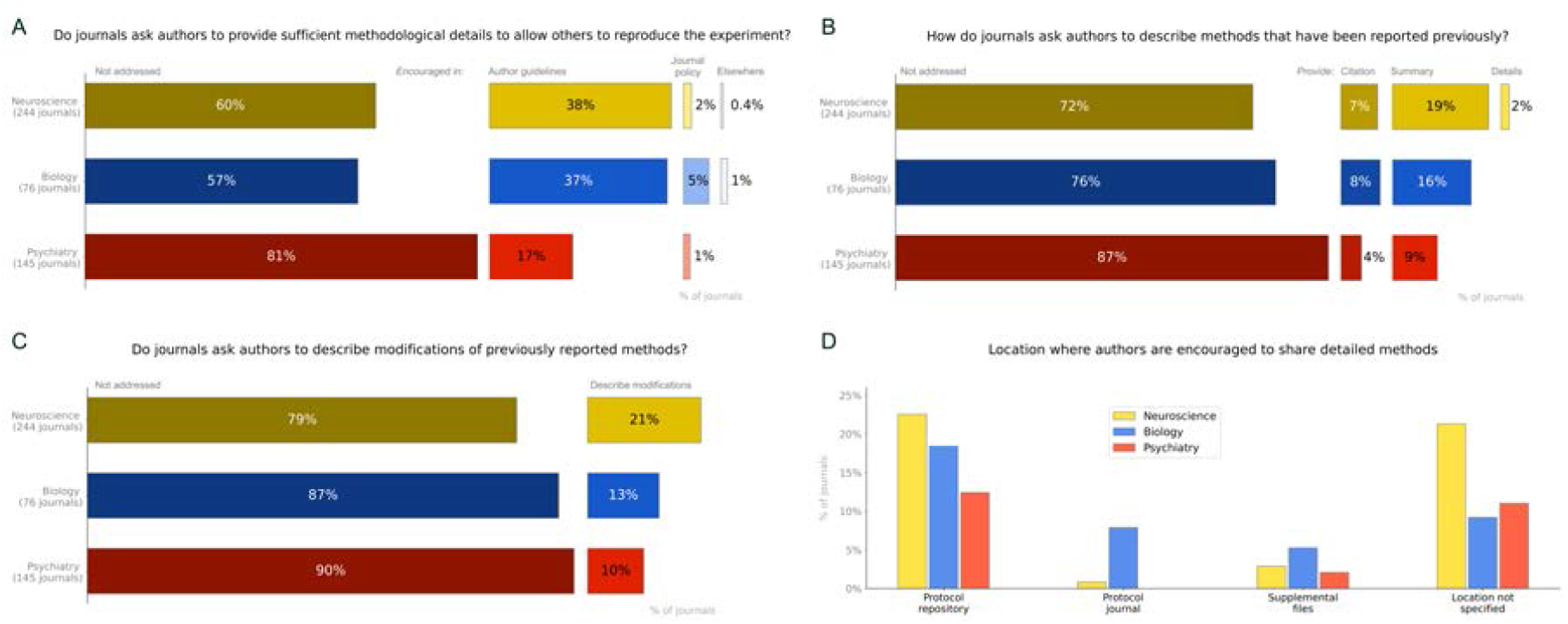
Journal policies for methodological reporting. Policies for all journals in three fields were assessed to determine (A) Whether journals ask authors to provide sufficient information about the methods to allow others to reproduce the experiments; (B) How journals ask authors to describe methods that have been reported previously; (C) Whether journals ask authors to describe modifications of previously reported methods; and (D) Where journals ask authors to share detailed methods. Percentages may not total 100% due to rounding errors.

### Reporting of previously described methods

Most journals had no policies concerning the reporting of methods that have been described previously (72-87%, Figure 5b). Some journals asked authors to summarize the method (9-19%) or to provide a citation instead of describing previously published methods in detail (4-8%). Policies asking authors to fully describe previously published methods were rare (0- 2%). Policies asking authors to report modifications of previously described methods were also uncommon (10-21%, Figure 5c).

### Where to share detailed methods

The percentage of journals that encouraged authors to share detailed methods in protocol repositories ranged between 12 and 23% (Figure 5d). A similar proportion of journals (9 to 21%) encouraged authors to share detailed methods elsewhere, without specifying where methods should be shared. Policies encouraging authors to share detailed methods in a protocol journal (0 to 8%) or as supplemental files (2 to 5%) were rare.

## Discussion

This exploratory study of papers in biology, neuroscience and psychiatry revealed several important findings. First, citations are often used in methods sections. Shortcut citations, explaining how a method was performed, were also routinely used. Other common reasons for using citations in the methods included “who or what” citations, that give credit to the authors of another paper or specify what was used, and “why” citations, which provide context or a justification. Different methods evolve at different rates, however citation age assessments suggested that some methods described in shortcut citations may no longer be state-of-the-art. The case series study showed that while shortcut citations can be used effectively, they can also create problems for readers seeking detailed methods. These problems included difficulty identifying the correct citation, accessing the cited materials, finding the cited method within the cited materials, and insufficient descriptions of the cited method. Following chains of shortcut citations to find methodological details was time-consuming, and each additional step in the chain can amplify the problems described in the previous sentence. Journals typically lack policies addressing methodological reporting. Fewer than one quarter of journals had policies addressing how authors should report methods that have been described previously, or address modifications of previously described methods. While some journals (18-44%, depending on the field) asked authors to provide sufficient methodological details to allow others to reproduce the method, most journals (57-81%) had no such policy.

### Using shortcut citations to foster a culture of open and reproducible methods

When using shortcut citations, authors replace a section of their methods with a citation referring to another resource. The details contained within that resource are essential to implement the method. We therefore propose that methodological shortcut citations should meet higher standards than other types of citations. Box 1 outlines three proposed criteria that authors can use to determine whether a resource should be cited as a shortcut. The open access criterion may be controversial for some, as it suggests that scientists who have a paywalled resource that meets the other two criteria should cite this resource to give credit, and create a second, open access resource (e.g. a deposited protocol) to cite as a shortcut. Nevertheless, we believe that the open access criteria is particularly important. Unlike other types of citations, readers who want to implement the study methods will need to read shortcut citations. Paywalled shortcut citations systematically deprive scientists of information needed to reproduce experiments.

#### Box 1: How to use methodological shortcut citations responsibly

Methodological shortcut citations replace a section of the methods. Detailed methods are essential for those seeking to implement the method. We therefore propose that resources cited as methodological shortcuts should meet higher standards than materials cited for other purposes.

We propose that authors should use three criteria to determine whether a paper, or another resource, should be used as a shortcut citation.

1. **Detailed description:** The resource should provide enough detail about the method that was used to allow others, including those who have little prior experience with the method, to implement the method. Resources with sufficient detail to be shortcut citations might include protocols, methods papers that are recent enough to reflect the current state of the art, or original research articles with unusually detailed methods.
2. **Similar or identical method:** The method described in the resource should be similar or identical to the method used by the authors. The authors should be able to describe any modifications to the methods in the methods section of their paper.
3. **Open access:** Paywalled or inaccessible shortcut citations deprive some readers of the information needed to implement the method. This creates disproportionate obstacles for researchers with limited access to publications, including researchers in countries with limited research funding and scientists who are not affiliated with a major institution. Paywalls can also limit access for researchers at well-funded centers. Researchers in Germany, for example, have not had access to Elsevier publications since January 1, 2018 due to ongoing subscription disputes [21]. Researchers within the University of California system lost access to Elsevier journals from 2019 to 2021 due to a similar subscription contract dispute [22].

Authors using shortcut citations should:

1. **Describe modifications in detail:** Deviations from the published method should be clearly described, in enough detail to allow others to implement the method. When a protocol posted on a repository is cited as a shortcut citation, the authors can version or fork the protocol to share their exact methods.
2. **Specify the exact location of the cited methods:** This might include providing page numbers where the cited method was described for books and manuals or describing the method name and location in the cited resource.

When a resource does not meet the criteria proposed in Box 1, we recommend that authors cite the resource to give credit to its authors and use other strategies to share detailed methods. Options for sharing detailed methods include supplemental files, protocol repositories and protocol journals (Table S7). Authors who deposit or publish protocols can cite these resources as shortcut citations.

Sharing methods in supplemental files is suboptimal for several reasons. First, readers who have access to the paper may not have access to the supplement. While completing this research, we noticed that some publishers and journals make supplemental files freely available, whereas others do not. Papers obtained through online repositories or interlibrary loan programs may not include supplements. Second, methods in supplemental files are not findable. While scientists can quickly search protocol repositories to identify relevant protocols, there is no way to identify papers that contain detailed methods in the supplemental files. Third, supplemental methods are often written as general descriptions, which are typically less useful than the detailed, step-by-step protocols shared in protocol repositories and methods journals. Fourth, supplemental methods cannot be updated after publication; hence, this format is not useful for sharing updated versions and tracking protocol evolution. This is especially important for methods that evolve rapidly.

A key advantage of protocol journals is that they allow authors to obtain credit for their methods development work through a peer-reviewed publication. One disadvantage is that the publication process takes time and effort. Furthermore, the published method can’t be updated – it only shows what one lab is doing at a single point in time. In contrast, protocol repositories allow authors to create living protocols by quickly sharing updated versions [23]. Some repositories also allow forking, in which scientists can post a modified version of a protocol posted by someone else [23]. Versions and forks are linked to the original protocol. This credits the original authors for their work, while allowing researchers to see how the protocol evolved. PLOS One offers a new “Lab protocol” publication [24] that combines the advantages of both formats. Lab protocol publications consist of a protocol, deposited on protocols.io [23], paired with a brief peer reviewed publication that provides context for the protocol. Authors can demonstrate that the protocol works by citing previous publications that used the protocol, or providing new data obtained using the protocol.

The decision tree shown in Figure 6 is designed to aid scientists in using shortcut citations responsibly, while reporting methods more transparently. Achieving these goals requires a shift in incentives – the scientific community needs to recognize and value protocols as a product of scientific work, on par with publications. While depositing methods may take more time, it also benefits authors in the long term. Depositing protocols in a repository that allows versioning and forking allows researchers to track changes as the protocol evolves and determine what version of a protocol was used for a particular publication. Furthermore, repositories provide long-term access to protocols even when researchers haven’t used the protocol in years, have moved to another lab or institution, or the person responsible for the protocol has left the lab. Detailed deposited protocols can also make it easier for new lab members to learn new protocols and expand their skills. Finally, other scientists can use and cite deposited protocols. This may make it easier to establish collaborations with others who are building on one’s methods or work as a community to identify previously unknown factors that affect protocol outcomes.

**Figure 6:**
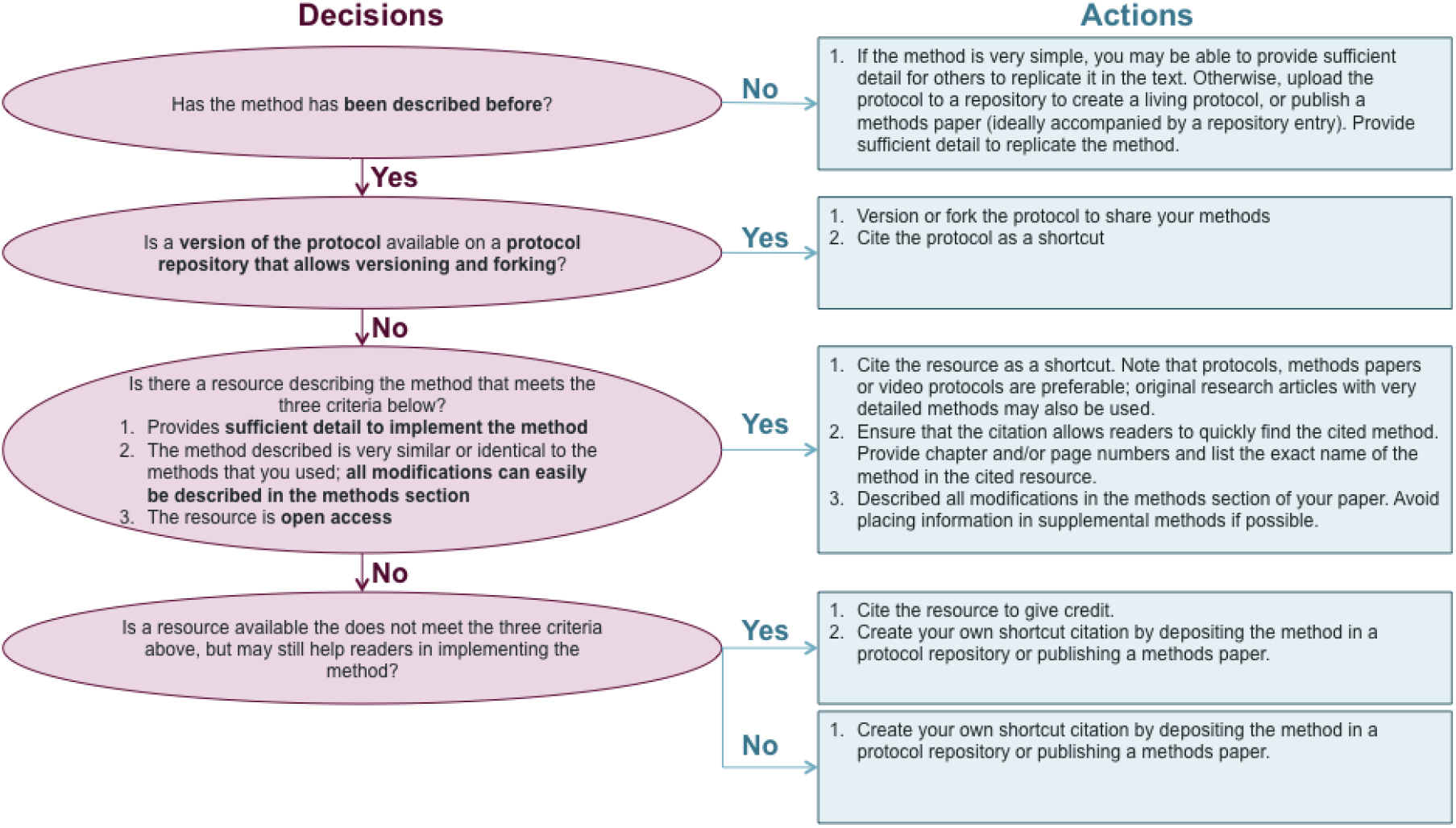
Decision tree for the responsible use of shortcut citations. This decision tree helps authors to prepare reproducible methods sections by determining when to use shortcut citations, and when to share detailed methods through protocol repositories or methods articles.

While some scientists believe that one should always cite the first paper to use a method, others prefer to cite more recent papers that fully describe and use methods that are similar to their own. These two beliefs reflect different reasons for citing a paper. Scientists who prefer to cite the original paper are using a “who or what” citation to give the authors of the cited paper credit for developing the method. In contrast, those who prefer to cite recent papers that use state-of-the-art protocols recognize that these citations are more useful as citation shortcuts. Fortunately, authors can do both. Authors should structure the citing sentence to help readers to distinguish between the “who or what” citation and the “shortcut” citation. For example, authors might write: *“Experiments were performed using an updated version [citation 1] of the protocol originally developed by Smith and colleagues [citation 2].”* This phrasing clearly demonstrates that the first citation is the shortcut citation, while the second citation is intended to give credit.

### The role of journals in fostering open and reproducible methods

The scientific community has rapidly expanding groups focused on open data and open code [25–28]; yet comparatively little attention has been dedicated to open methods. Open data are most useful when they are generated through rigorous experimental designs, using reproducible methods. Data are of little value when we don’t understand or can’t evaluate the quality of the methods used to generate them. By implementing the actions outlined below, journals can foster a culture of protocol sharing, and open and reproducible methods, within the scientific communities that they serve.

- **Make all methods publications open access.** Eliminating paywalls for methods papers would allow everyone to access the information needed to implement the methods described.
- **Replace static methods papers with dynamic formats where authors publish a living protocol on an open access protocol repository:** The static formats used to publish traditional research papers, which are primarily focused on results, do not work well for methods. Protocols evolve over time. The question is not whether protocols will change; but when and how they will change. Furthermore, step by step protocols are often more useful that the text descriptions found in methods section. Journals should embrace dynamic formats for publishing methods, such as the PLOS One Lab Protocols article [24]. Authors publish a detailed living protocol on an open access protocol repository [23], which can be versioned and forked to track the evolution of methods within and across research teams.
- **Ensure that the protocol repositories used to publish dynamic methods allow versioning and forking:** Versioning and forking allow readers to share and track the evolution of methods over time, both within and across labs. Forking benefits protocol creators by giving them credit for their work, while allowing them to see how others are adapting their methods. Without forking, researchers may be reluctant to share adapted versions of protocols created by others, as they do not want to inadvertently claim credit for work that was not their own.
- **Exempt methods sections from word count limits. Ask authors to provide sufficient details to allow others to implement the method.** Word count exemption policies [29] eliminate a barrier to sharing detailed methods.
- **Encourage authors to share methods in protocol repositories, not as supplemental files.** Protocols deposited in repositories are findable and dynamic, whereas methods deposited in supplemental files are not.
- **Require authors use methodological citation shortcuts responsibly, according to the criteria described in Box 1. Eliminate policies requiring authors to cite previous publications instead of fully describing methods.** The scientific community needs to rethink the way that shortcut citations are used to ensure that these citations contribute to a culture of open and reproducible methods reporting. Policies that ask authors to use shortcut citations irresponsibly deprive others of details needed to implement the method. The driving force behind some of these policies may be concerns about plagiarism and copyright violations [5]. Laws in some countries specifically exempt detailed descriptions of protocols and methods from copyright restrictions (e.g. United States [30, 31], European Union [32]), whereas other countries take a more indirect approach by not including detailed descriptions of methods under items that can be copyrighted (e.g. United Kingdom [33, 34]). Other countries may not offer similar protection. Resolving these legal issues is crucial to allow scientists to fully describe their methods whenever they publish their work.

### Limitations

This study has several important limitations. Data examining the reasons for citing a paper in the methods (Figure 2a) should be viewed as a rough approximation. Abstractors were making distinctions about the reasons for a citation that the authors themselves likely did not make when inserting citations into their methods sections. Small changes in the wording or position of the reference could alter the categorization, as could variations in reader expertise. When abstractors encountered a citation that could potentially fall under multiple categories, they were instructed to choose the most likely category. Abstractors encountered many citations that could have been classified under more than one category. Researchers who use this protocol in the future may wish to allow abstractors to select more than one category for each citation. Probable and possible shortcuts were defined using syntactic definitions. Conceptual definitions were not feasible, as reviewers’ familiarity with the methods described was highly variable. Our results may not apply to journals with lower impact factors, or non-English language journals. The case series was designed to determine what problems one might encounter when following shortcut citations in search of detailed methods. Data from this small, non-representative sample (1 paper per quintile of possible plus probable shortcut citations per field) should not be used to determine how often each of these problems occurred. Larger samples would be needed to answer this question. This study focused on shortcut citations. We did not examine the completeness of methodological reporting when authors described a method without using a shortcut citation.

## Conclusions

Many authors used methodological shortcut citations to explain their research methods. While shortcut citations can be used effectively, they can also make it difficult for readers to find methodological details that would be needed to implement the method. Journals often lack clear policies to encourage open and reproducible methods reporting. We propose that methodological shortcut citations should meet three criteria. Cited resources should describe a method similar to the one used by the authors, provide enough detail to allow others to implement the method, and be open access. We outline additional steps that authors can take to use methodological shortcut citations responsibly, and also propose actions that journals can take to foster a culture of open and reproducible methods reporting. Future studies should explore strategies for creating and sharing detailed protocols that others can use. The scientific community should also identify opportunities to incentivize and reward open methods.

## Funding

This study was completed through a participant guided, learn-by doing meta-research course funded by the Berlin University Alliance within the Excellence Strategy of the federal and state governments (301_TrainIndik).

## Acknowledgements

The authors thank Małgorzata Anna Gazda and Alberto Antonietti for their assistance in translating policies of Polish and Italian journals.

## CREDIT Authorship Statement

### Conceptualization

Tracey Weissgerber, Kathleen Bastian, Britta Lewke, Orestis Rakitzis, Anna-Delia Knipper, Philipp van Kronenberg Till

### Methodology

Tracey Weissgerber, Kathleen Bastian, Britta Lewke, Orestis Rakitzis, Anna-Delia Knipper, Philipp van Kronenberg Till, Vartan Kazezian

### Investigation

Vartan Kazezian, Anna-Delia Knipper, Shambhavi Chidambaram, Philipp van Kronenberg Till, Kathleen Bastian, Bruna Seco, Nina Nitzan Soto, Kai Standvoss, Fereshteh Zarebidaki, Subhi Arafat, Britta Lewke, Ana Herrera-Melendez, Isa Steinecker, Orestis Rakitzis, Ubai Alsharif

### Project administration

Tracey Weissgerber, Vartan Kazezian, Britta Lewke, Kai Standvoss

### Data curation

Kai Standvoss, Vartan Kazezian, Isa Steinecker

### Formal analysis

Kai Standvoss, Nina Nitzan Soto

### Software

Britta Lewke

### Supervision

Tracey Weissgerber, Vartan Kazezian

### Validation

Kai Standvoss, Britta Lewke, Vartan Kazezian, Isa Steinecker, Tracey Weissgerber

### Visualization

Kai Standvoss, Tracey Weissgerber, Nina Nitzan Soto, Vartan Kazezian, Shambhavi Chidambaram, Isa Steinecker, Ana Herrera-Melendez, Fereshteh Zarebidaki, Anna-Delia Knipper, Orestis Rakitzis, Philipp van Kronenberg Till

### Writing original draft

Tracey Weissgerber, Shambhavi Chidambaram, Kathleen Bastian, Anna-Delia Knipper, Isa Steinecker, Bruna Seco, Vartan Kazezian, Britta Lewke

### Writing – reviewing and editing

Vartan Kazezian, Anna-Delia Knipper, Shambhavi Chidambaram, Philipp van Kronenberg Till, Kathleen Bastian, Bruna Seco, Nina Nitzan Soto, Kai Stondvoss, Fereshteh Zarebidaki, Subhi Arafat, Britta Lewke, Ana Herrera-Melendez, Isa Steinecker, Orestis Rakitzis, Ubai Alsharif, Tracey Weissgerber

### Funding

Tracey Weissgerber

## Supplementary Materials Legend

**Figure S1:**
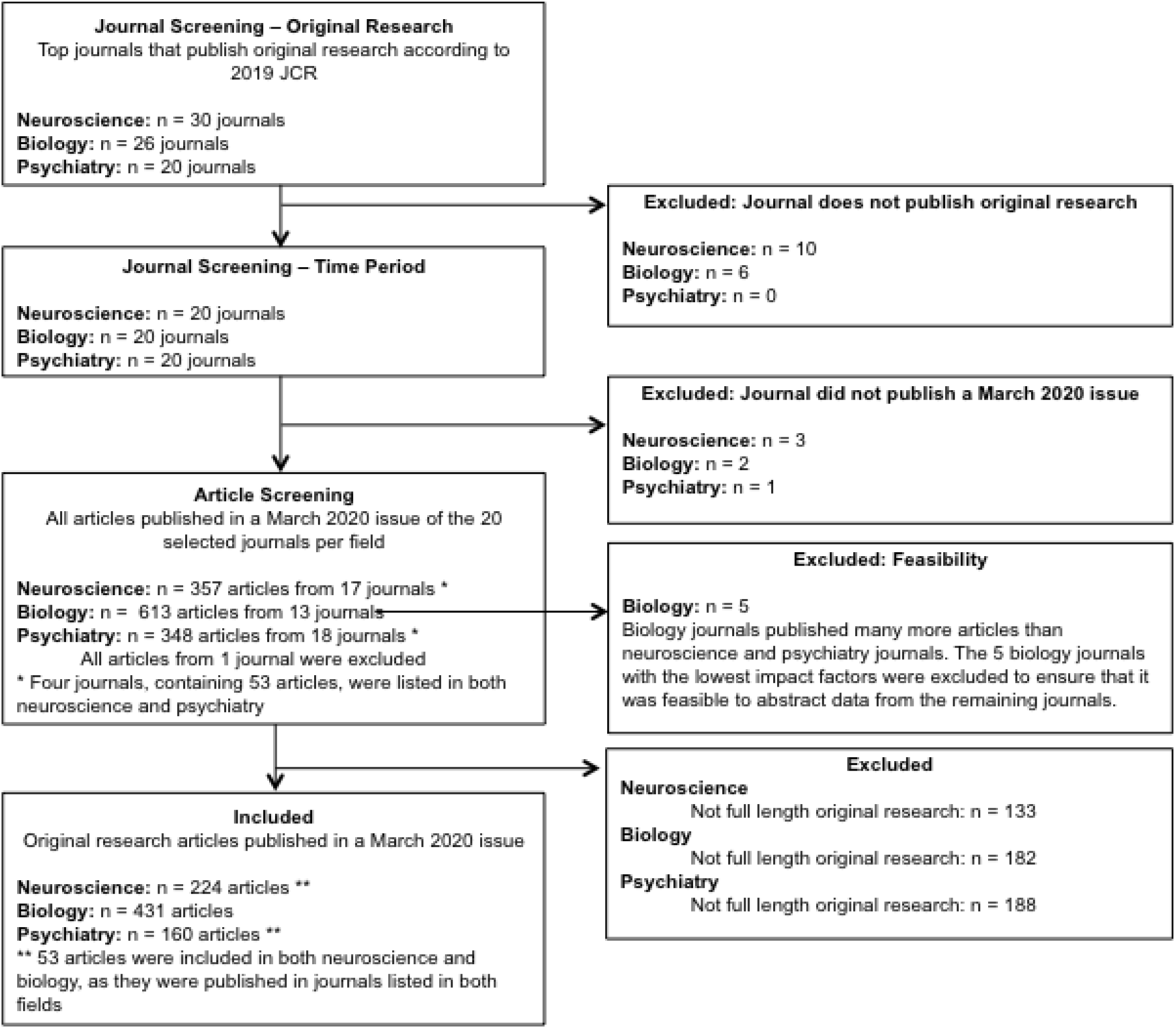
Prevalence study flow chart. This flow chart illustrates the journal and article screening process and shows the number of observations excluded and reasons for exclusion at each phase of screening.

**Figure S2:**
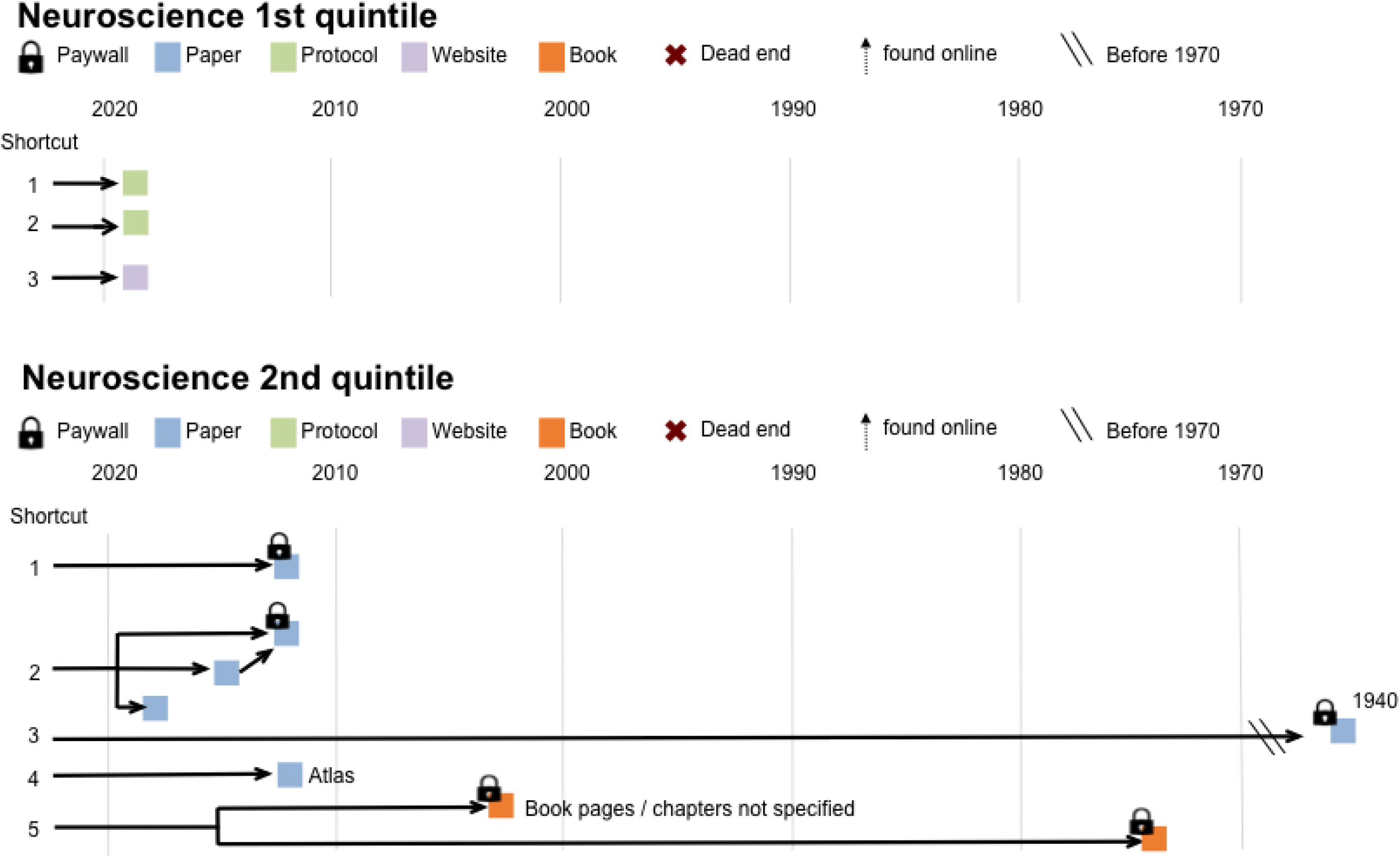
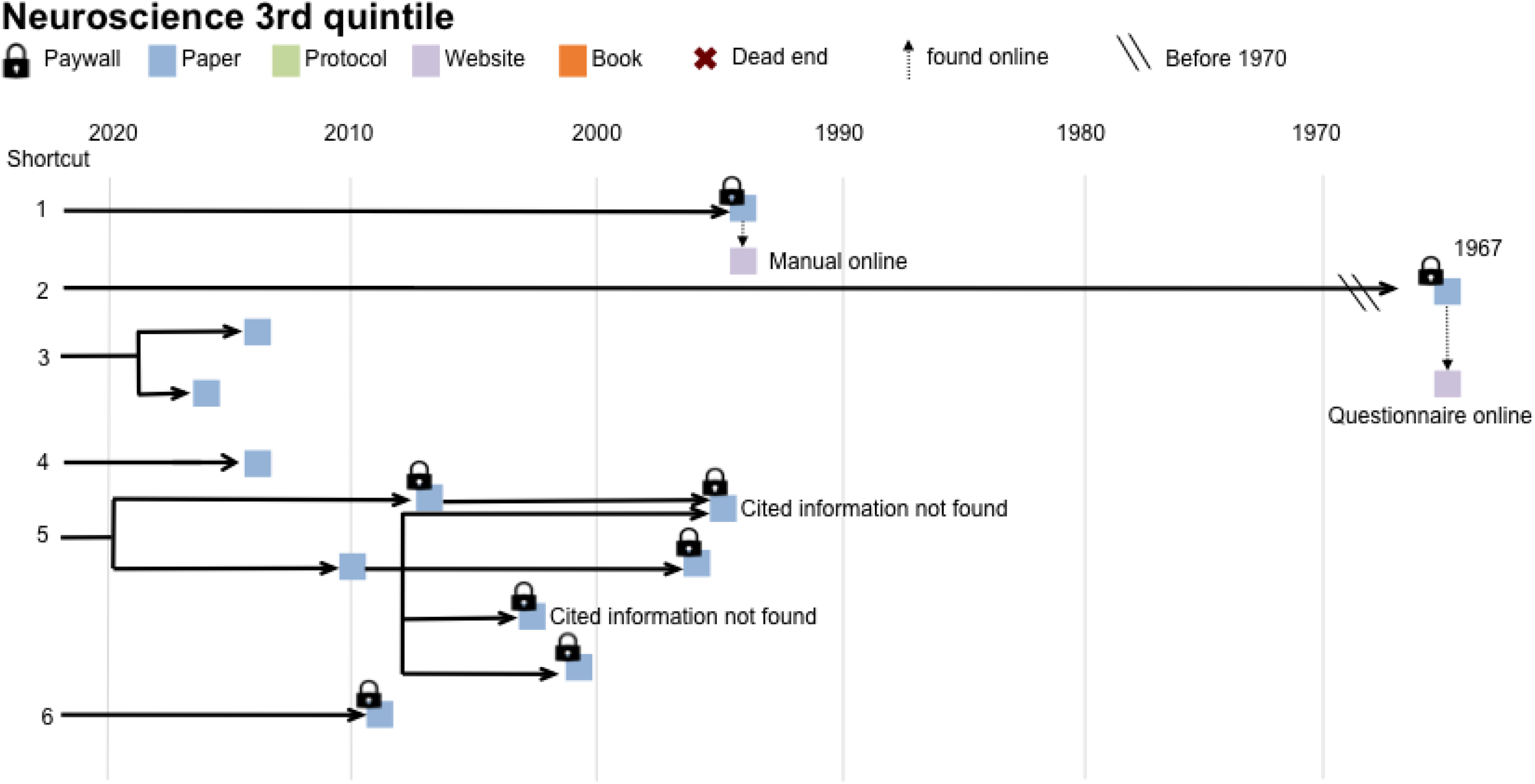
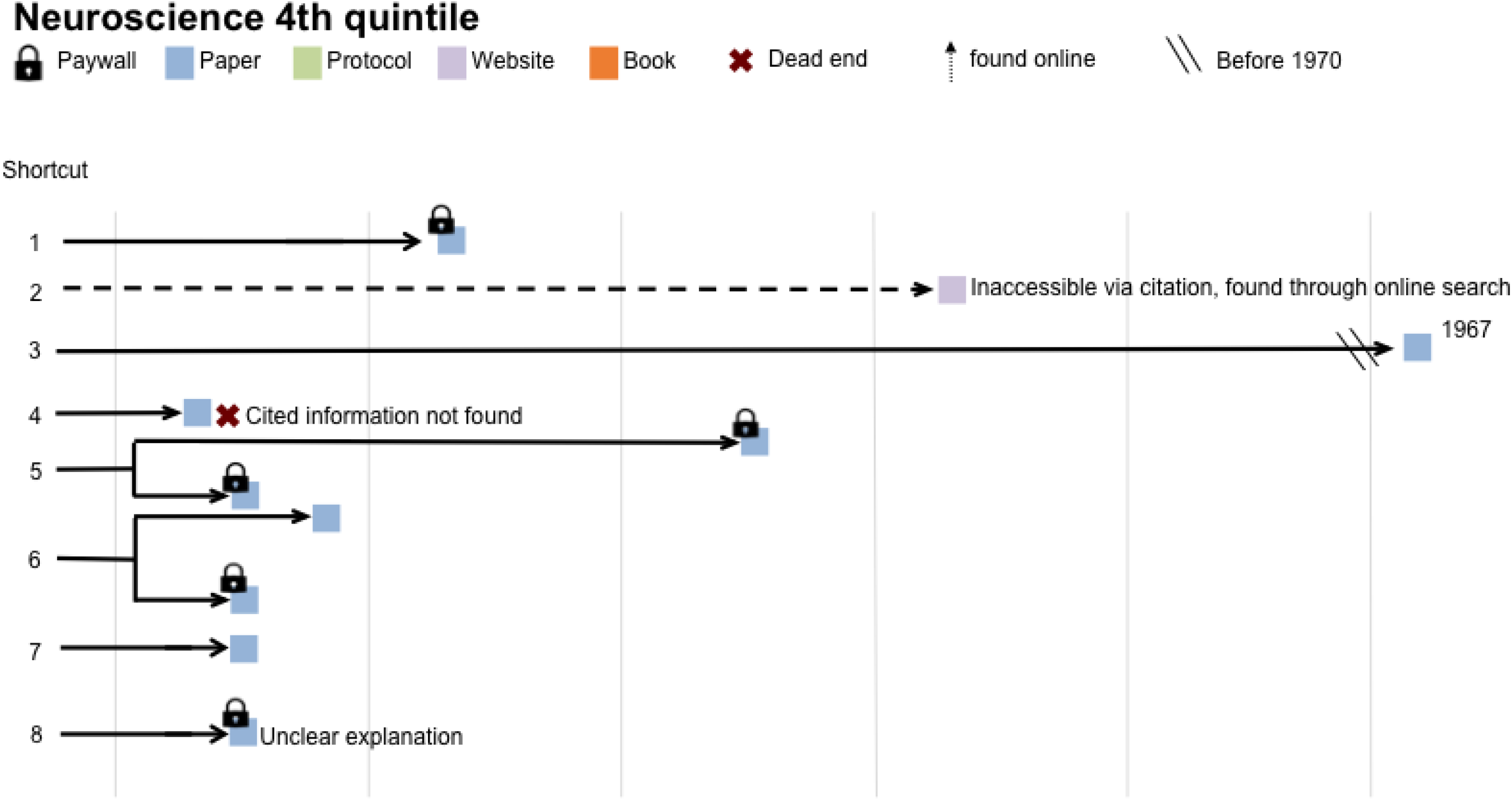
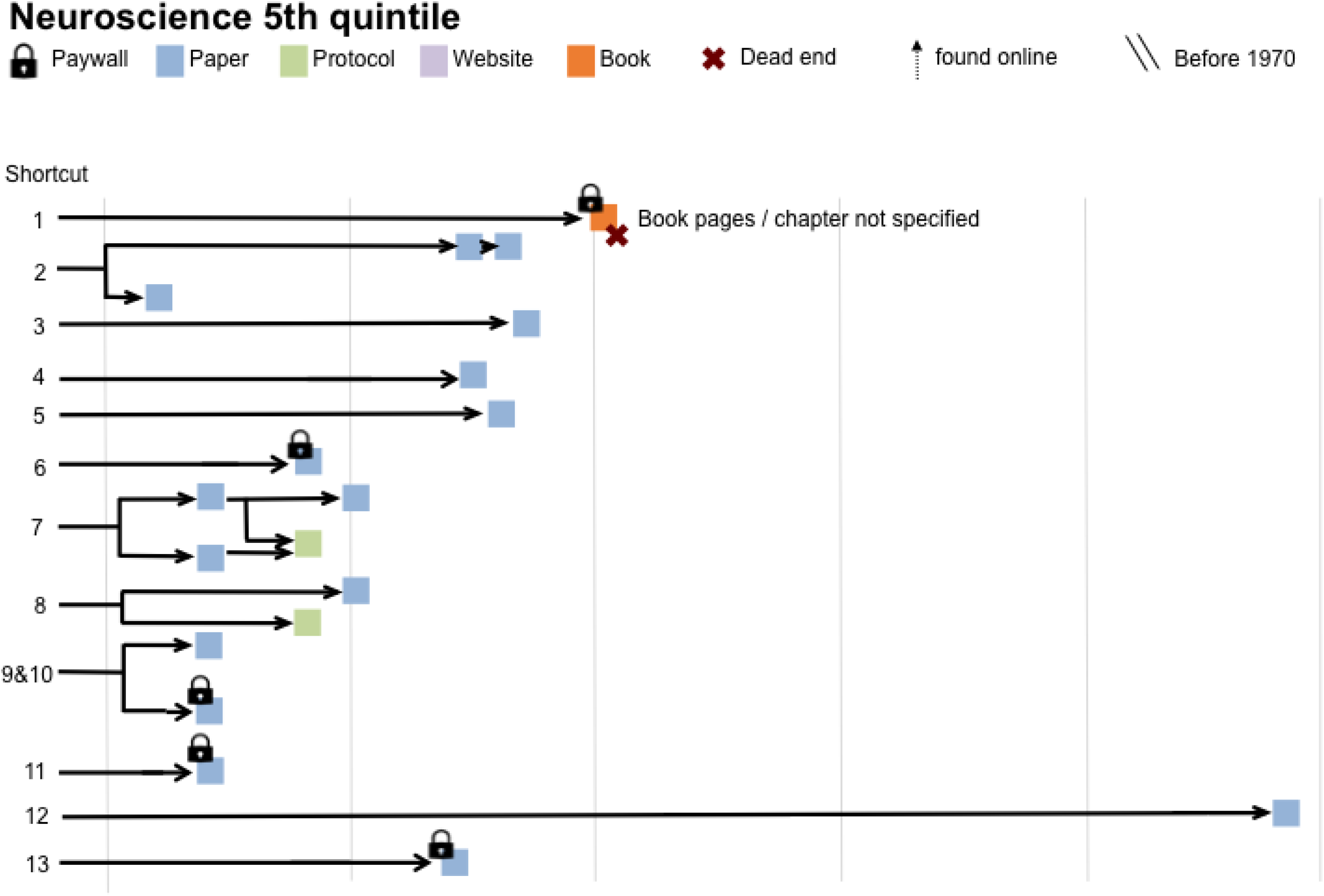
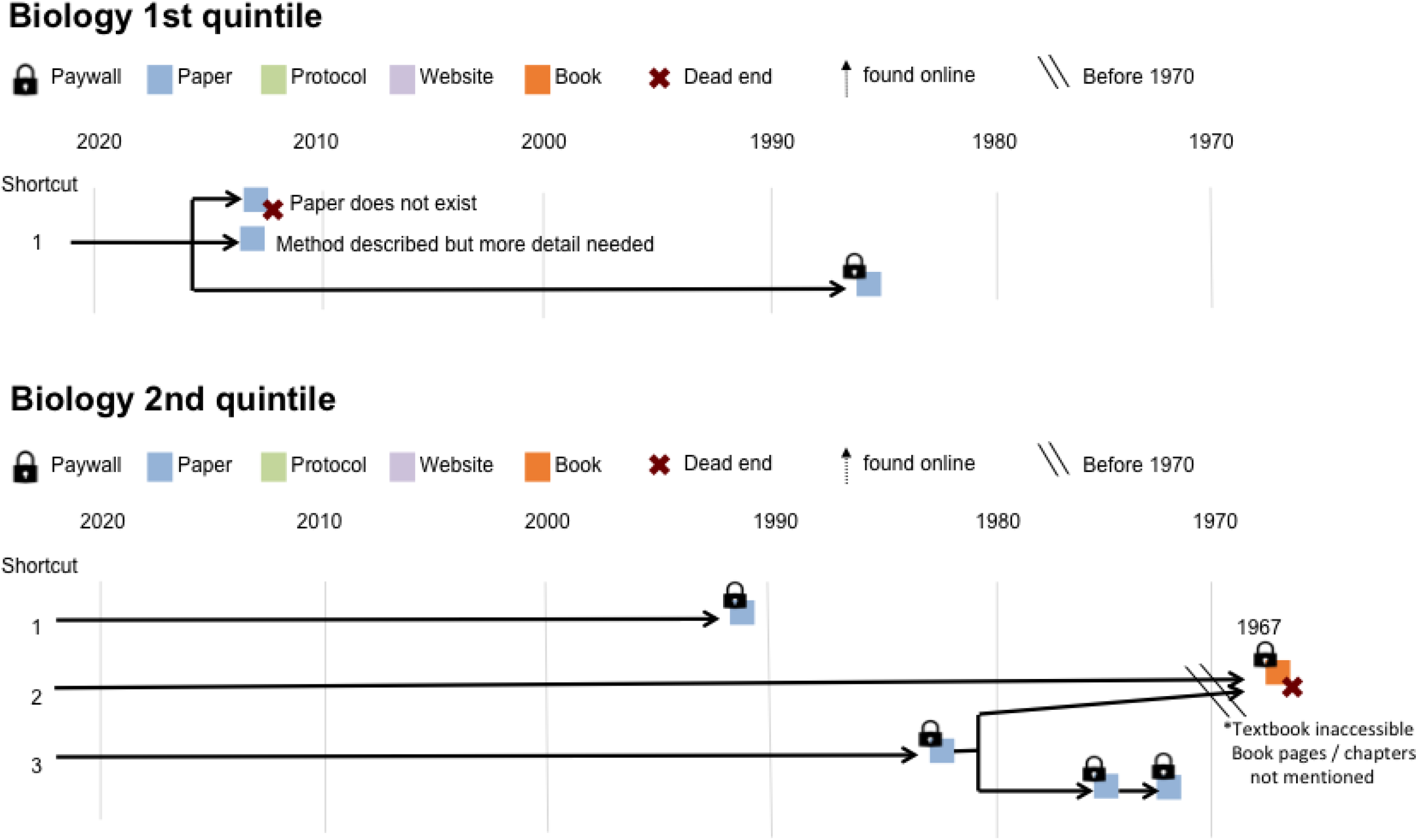
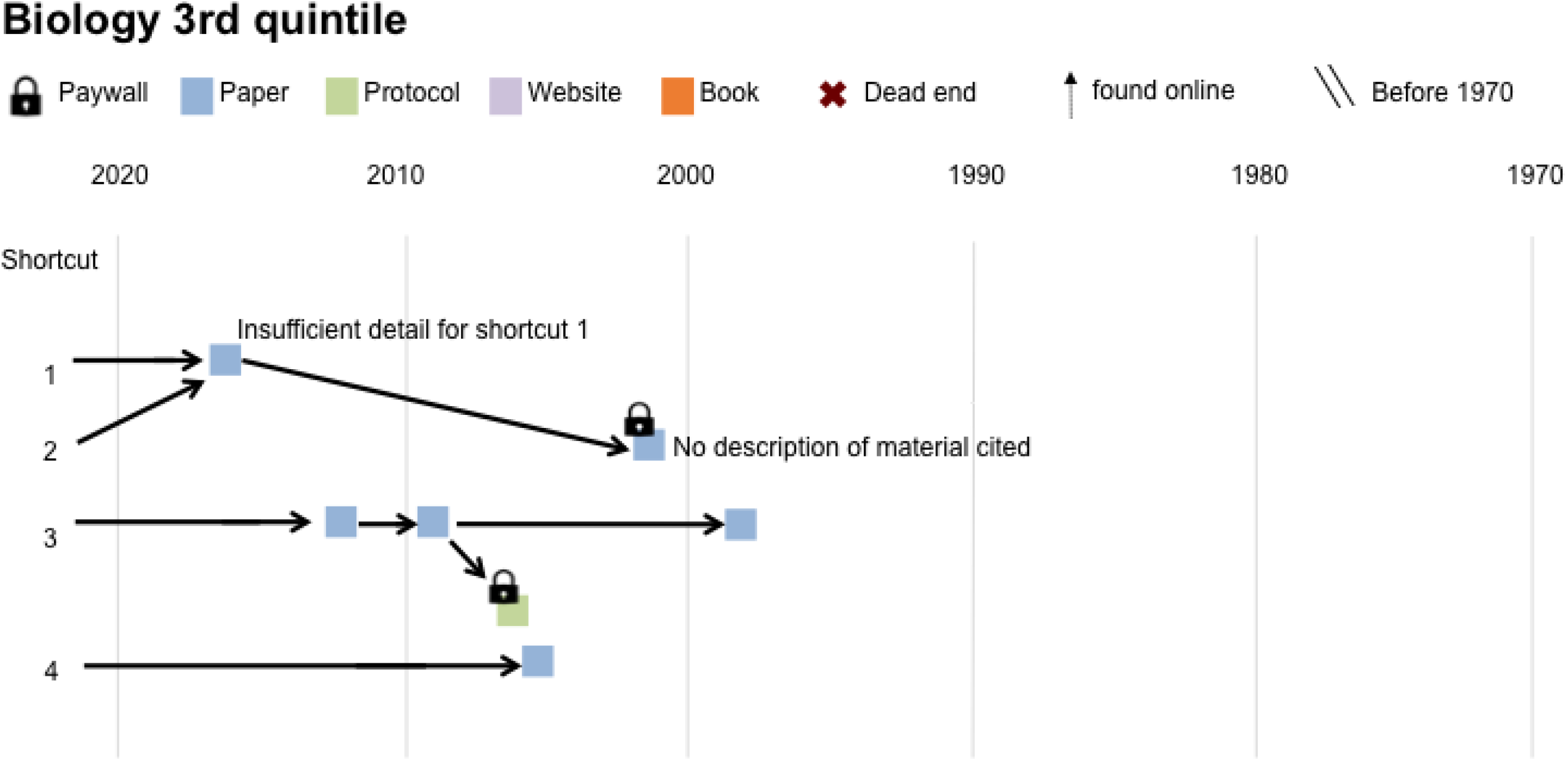
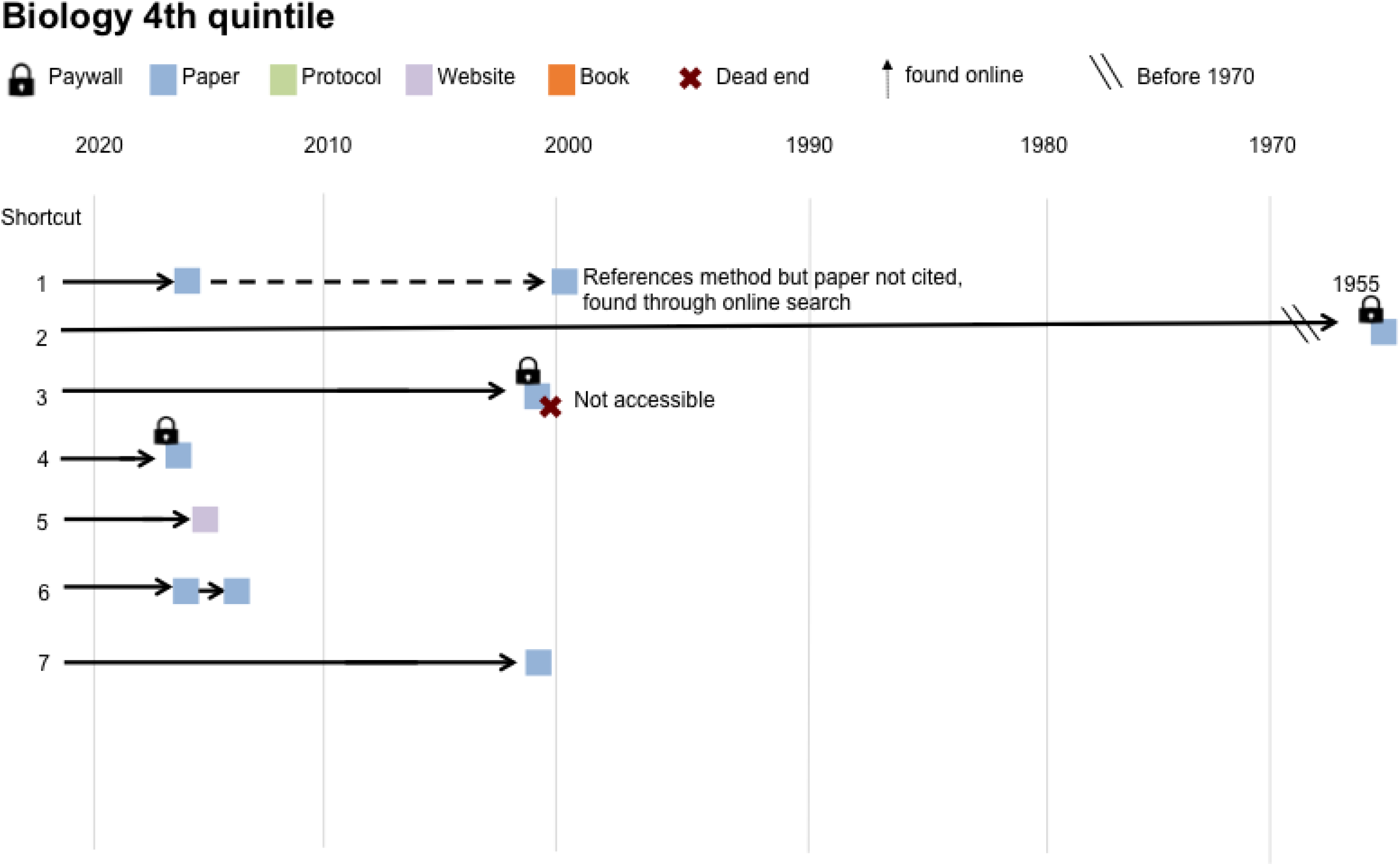
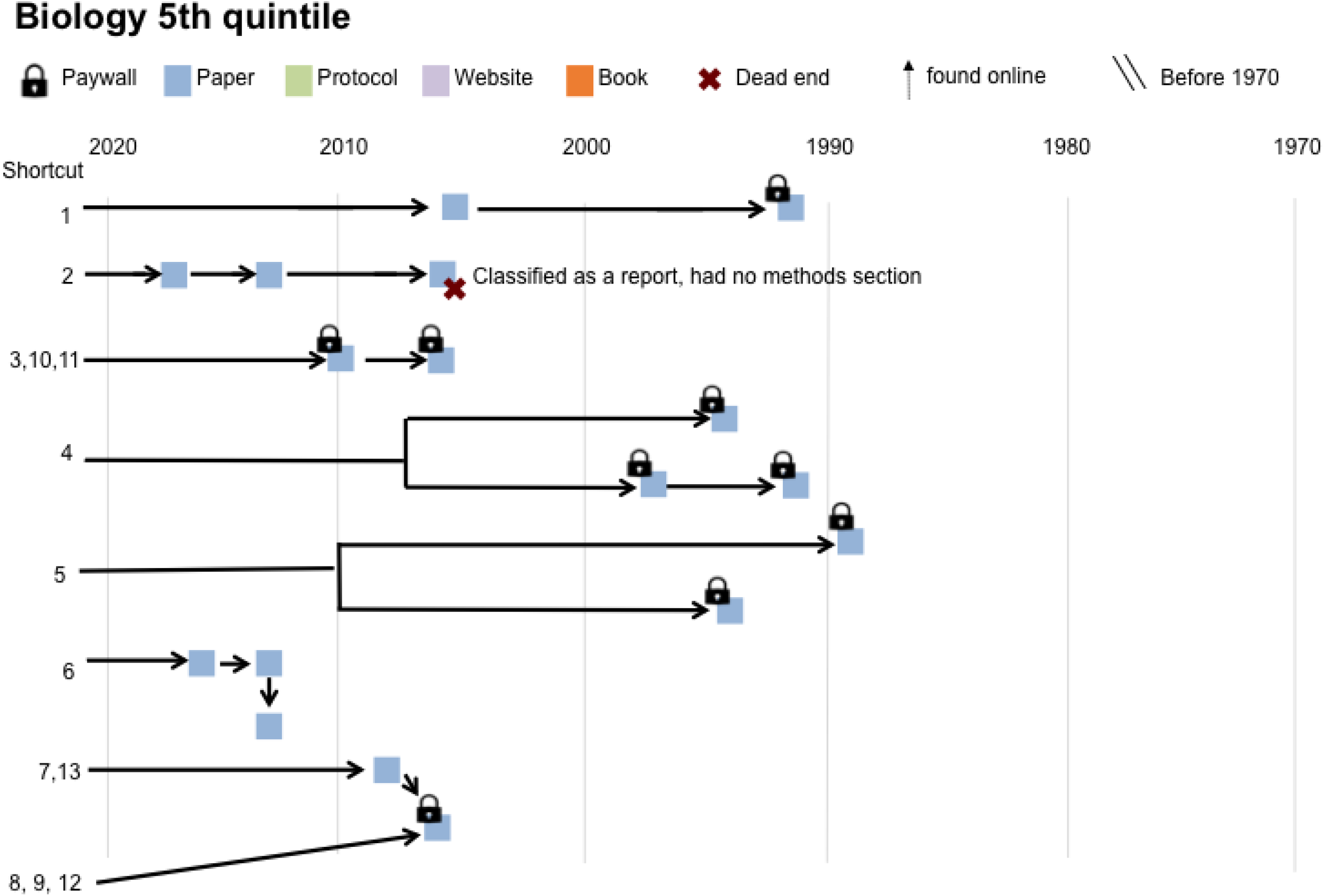
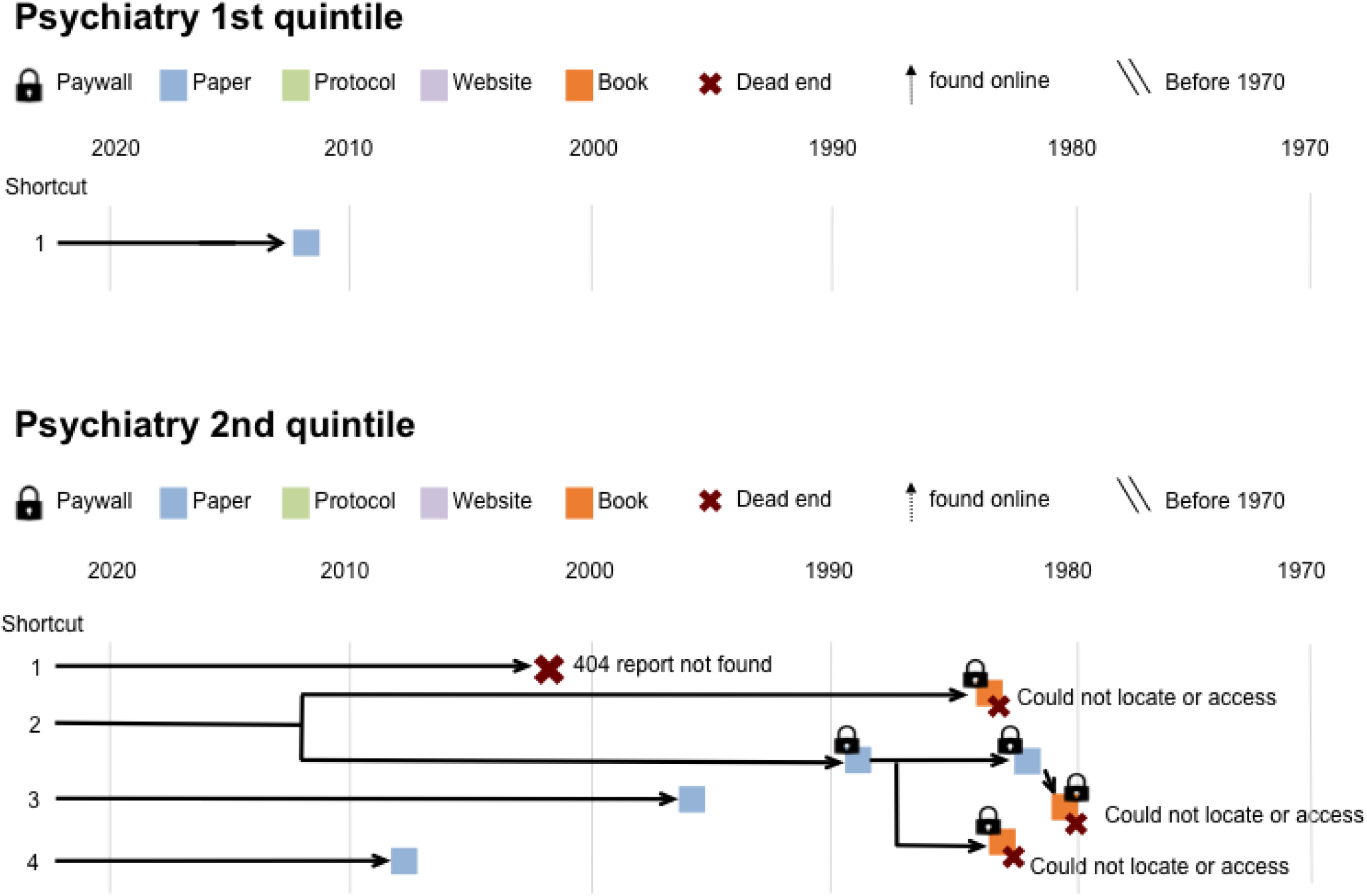
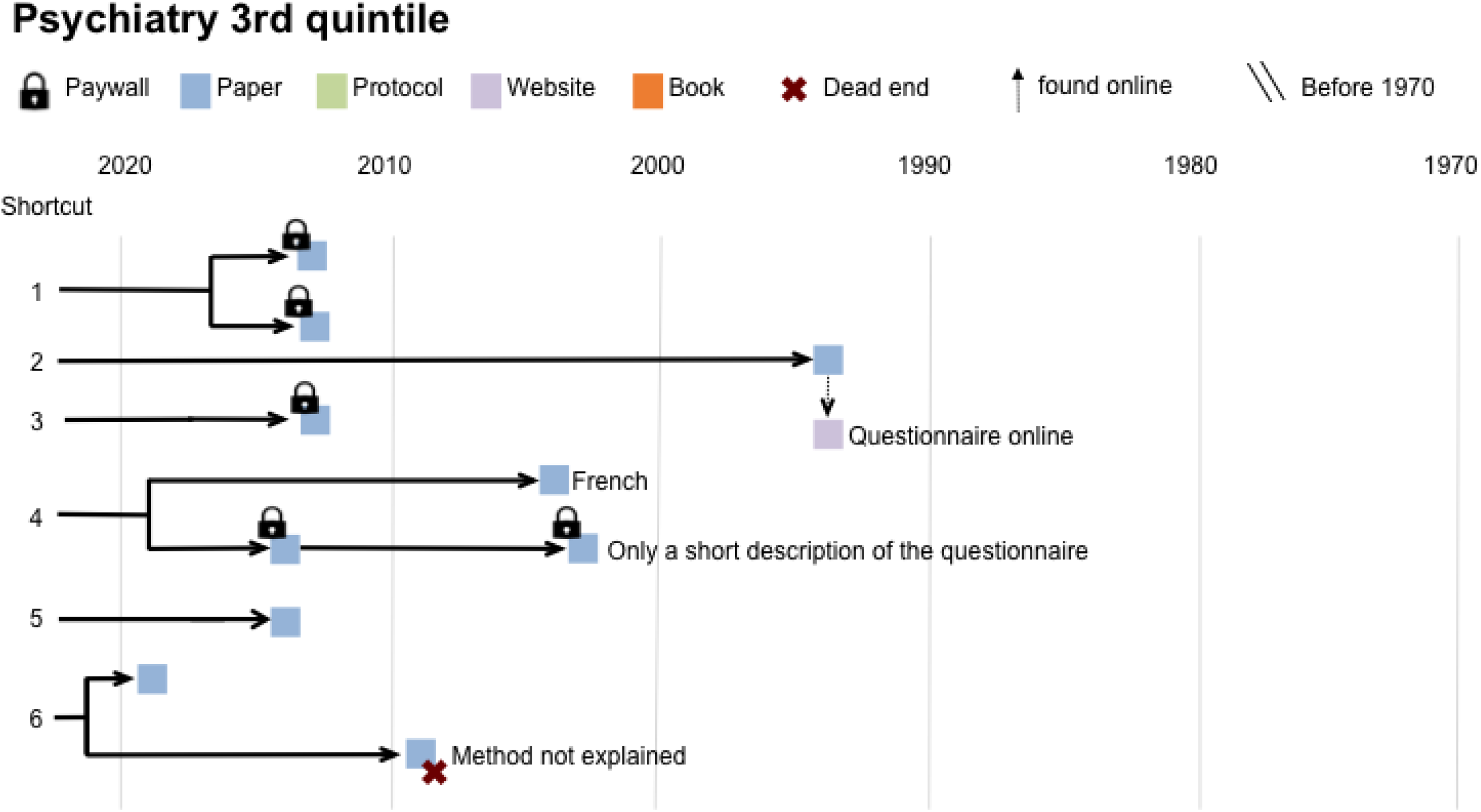
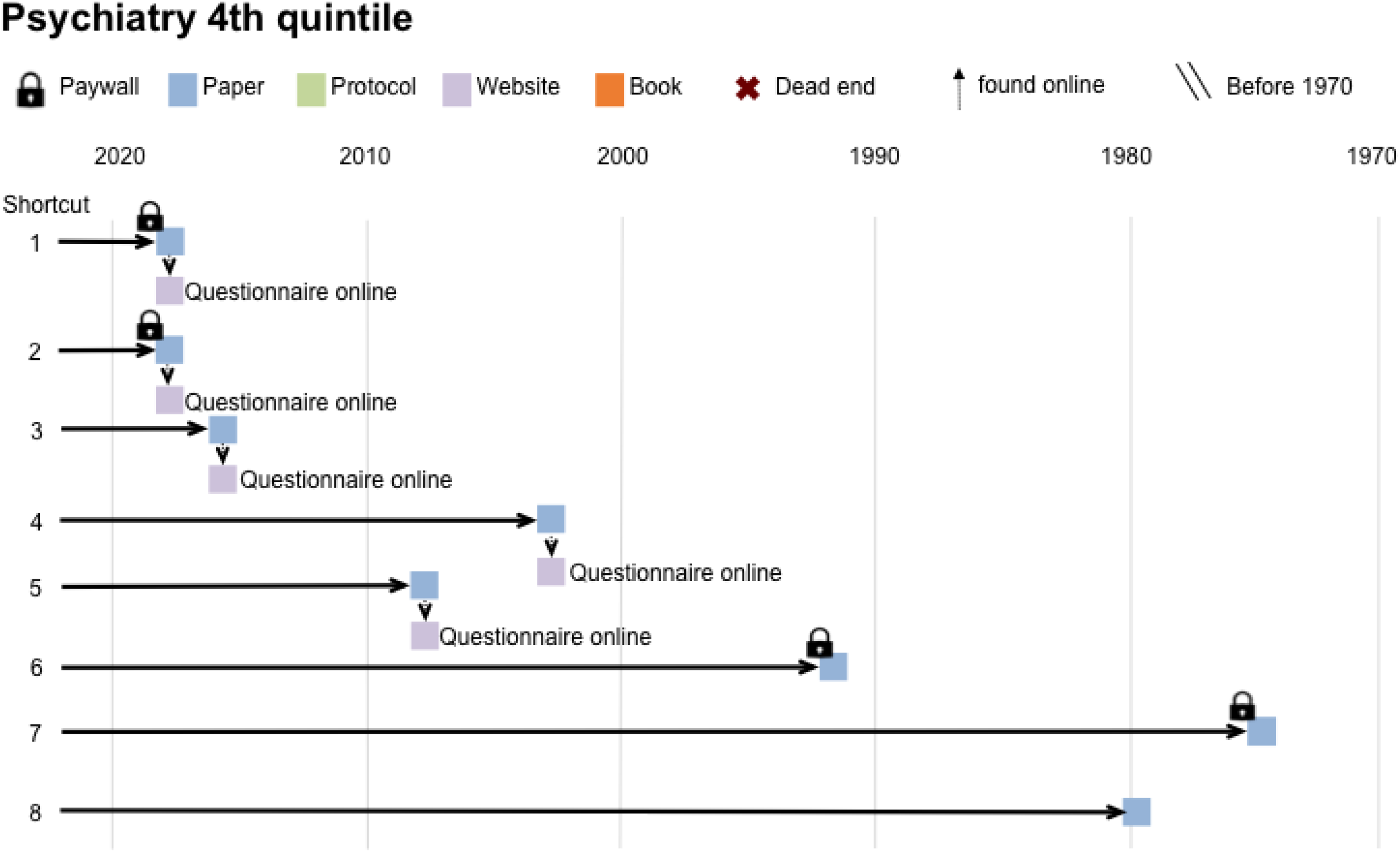
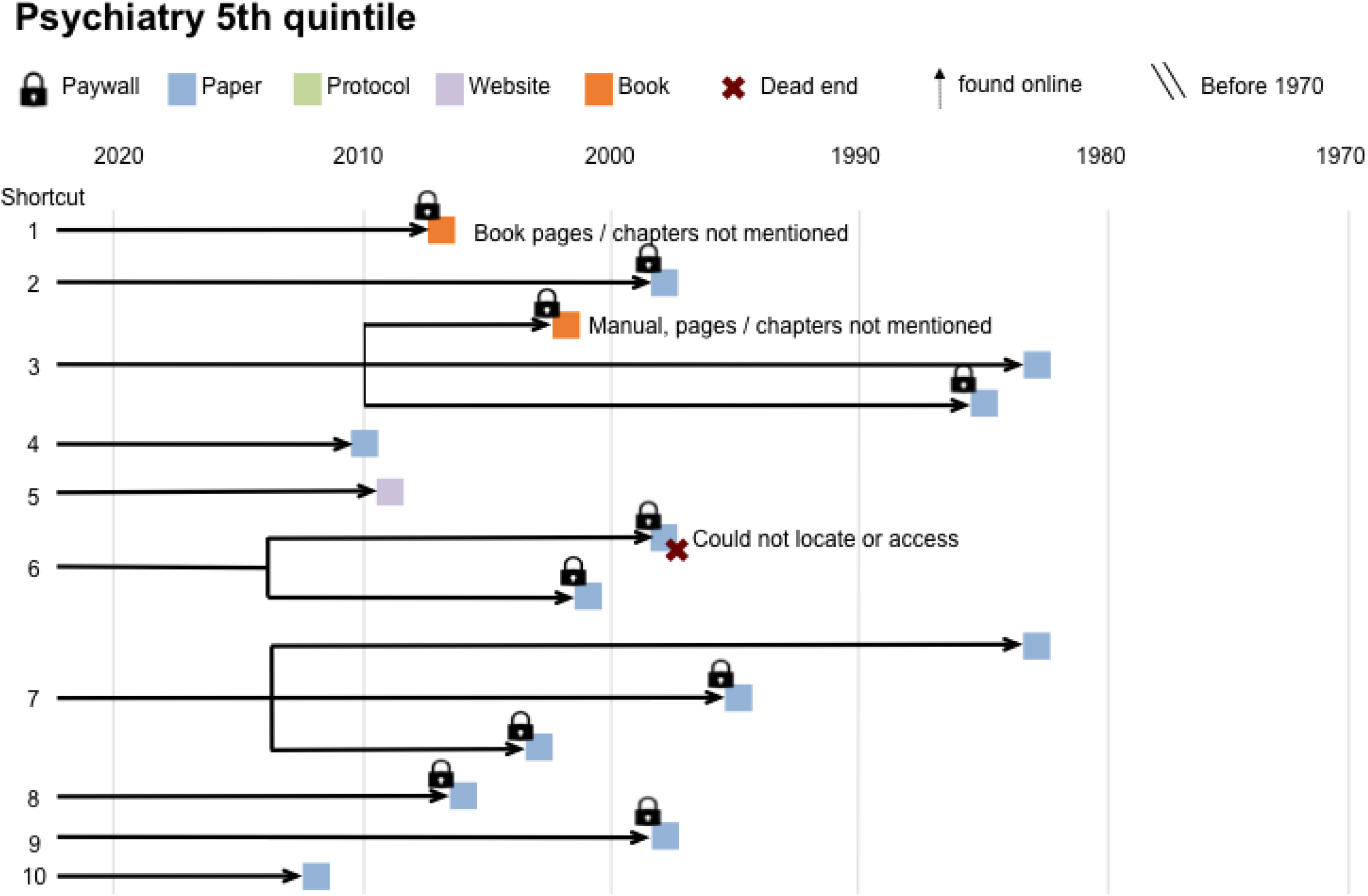
Case series diagrams for individual articles. These diagrams map the process of finding methodological details for each of the 15 papers in the case series. Reviewers consulted resources cited in shortcut citations to find methodological details. The diagrams show the publication year and type of each cited resource and whether the resource was behind a paywall. Chains of shortcut citations occur when the cited source also uses a methodological shortcut citation to describe the method. Text on the diagram provides information describes problems encountered when searching for details about the cited method.

**Figure S3:**
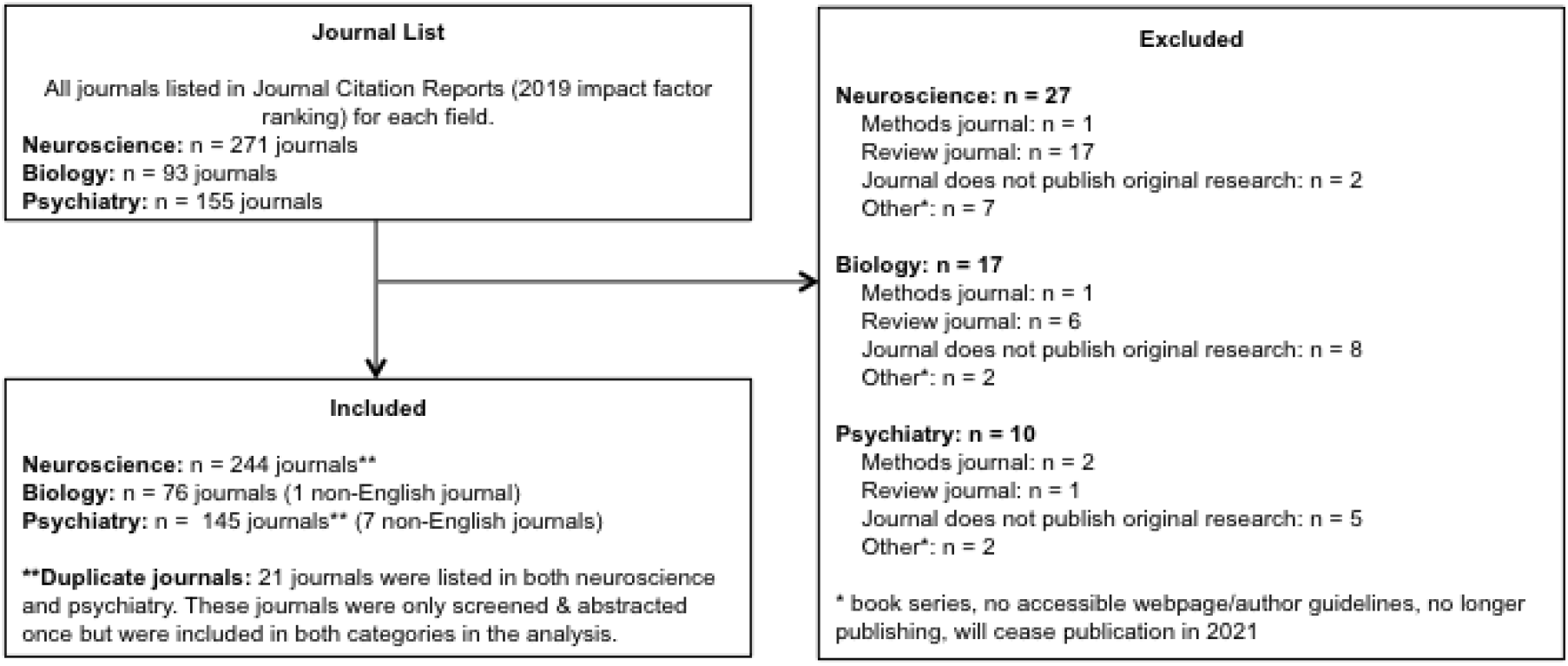
Flow diagram for journal policy study. This flow chart illustrates the journal screening process and shows the number of obsevations excluded and reasons for exclusion at each phase of screening.

**Table S1:**
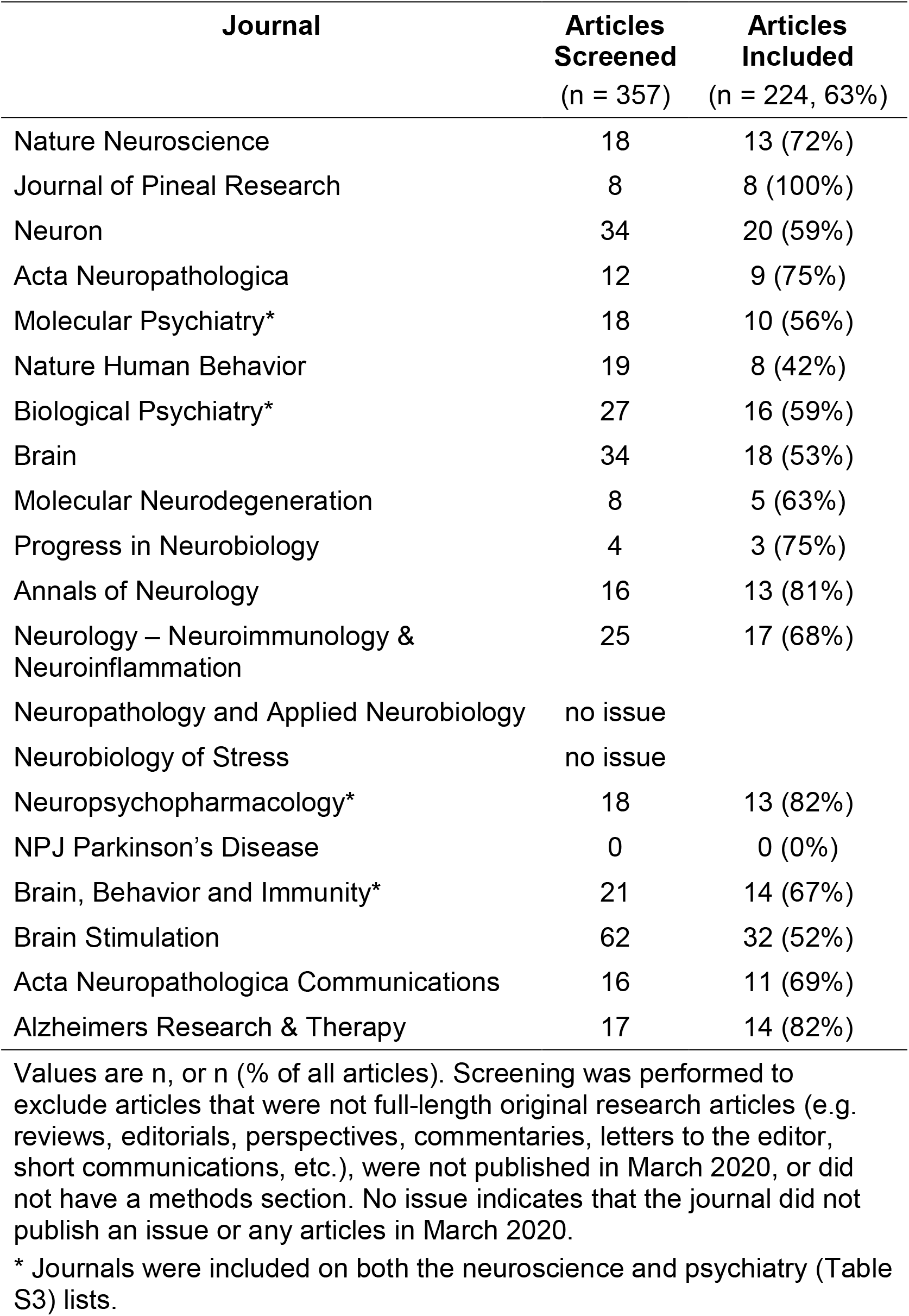
Number of articles examined for each neuroscience journal.

| **Journal** | **Articles**<br>**Screened**<br>(n = 357) | **Articles**<br>**Included**<br>(n = 224, 63%) |
| --- | --- | --- |
| Nature Neuroscience | 18 | 13 (72%) |
| Journal of Pineal Research | 8 | 8 (100%) |
| Neuron | 34 | 20 (59%) |
| Acta Neuropathologica | 12 | 9 (75%) |
| Molecular Psychiatry* | 18 | 10 (56%) |
| Nature Human Behavior | 19 | 8 (42%) |
| Biological Psychiatry* | 27 | 16 (59%) |
| Brain | 34 | 18 (53%) |
| Molecular Neurodegeneration | 8 | 5 (63%) |
| Progress in Neurobiology | 4 | 3 (75%) |
| Annals of Neurology | 16 | 13 (81%) |
| Neurology – Neuroimmunology & Neuroinflammation | 25 | 17 (68%) |
| Neuropathology and Applied Neurobiology | no issue |  |
| Neurobiology of Stress | no issue |  |
| Neuropsychopharmacology* | 18 | 13 (82%) |
| NPJ Parkinson's Disease | 0 | 0 (0%) |
| Brain, Behavior and Immunity* | 21 | 14 (67%) |
| Brain Stimulation | 62 | 32 (52%) |
| Acta Neuropathologica Communications | 16 | 11 (69%) |
| Alzheimers Research & Therapy | 17 | 14 (82%) |

Values are n, or n (% of all articles). Screening was performed to exclude articles that were not full-length original research articles (e.g. reviews, editorials, perspectives, commentaries, letters to the editor, short communications, etc.), were not published in March 2020, or did not have a methods section. No issue indicates that the journal did not publish an issue or any articles in March 2020.
\* Journals were included on both the neuroscience and psychiatry (Table S3) lists.

**Table S2:** Number of articles examined for each biology journal.

| **Journal** | **Articles**<br>**Screened**<br>(n = 613) | **Articles**<br>**Included**<br>(n = 431, 70%) |
| --- | --- | --- |
| Current Biology | 70 | 23 (33%) |
| eLife | 165 | 129 (78%) |
| PLoS Biology | 44 | 28 (64%) |
| BMC Biology | 16 | 13 (81%) |
| Philosophical Transactions of the Royal Society B - Biological Sciences | 52 | 24 (46%) |
| FASEB Journal | 89 | 82 (92%) |
| Bioelectrochemistry | no issue |  |
| Proceedings of the Royal Society B- Biological Sciences | 45 | 40 (89%) |
| Science China – Life Sciences | 17 | 8 (47%) |
| Geobiology | 7 | 7 (100%) |
| Communications Biology | 56 | 51 (91%) |
| Astrobiology | 8 | 8 (100%) |
| Biology | 22 | 15 (68%) |
| Yale Journal of Biology and Medicine | 22 | 3 (14%) |
| Interface Focus | no issue |  |

Values are n, or n (% of all articles). Screening was performed to exclude articles that were not full-length original research articles (e.g. reviews, editorials, perspectives, commentaries, letters to the editor, short communications, etc.), were not published in March 2020, or did not have a methods section. No issue indicates that the journal did not publish an issue or any articles in March 2020.

**Table S3:** Number of articles examined for each psychiatry journal.

| **Journal** | **Articles**<br>**Screened**<br>(n = 348) | **Articles**<br>**Included**<br>(n = 160, 46%) |
| --- | --- | --- |
| World Psychiatry | no issue |  |
| JAMA Psychiatry | 10 | 7 (70%) |
| Lancet Psychiatry | 31 | 3 (10%) |
| Psychotherapy and Psychosomatics | 13 | 1 (8%) |
| American Journal of Psychiatry | 17 | 5 (29%) |
| Molecular Psychiatry* | 18 | 10 (56%) |
| Biological Psychiatry* | 27 | 16 (59%) |
| Journal of Neurology, Neurosurgery and Psychiatry | 21 | 9 (43%) |
| Schizophrenia Bulletin | 28 | 17 (61%) |
| British Journal of Psychiatry | 16 | 5 (31%) |
| Journal of Child Psychology and Psychiatry | 19 | 0 (0%) |
| Journal of the American Academy of Child and Adolescent Psychiatry | 18 | 9 (50%) |
| Neuropsychopharmacology* | 18 | 13 (72%) |
| Brain, Behavior and Immunity* | 21 | 14 (67%) |
| Addiction | 29 | 14 (48%) |
| Epidemiology and Psychiatric Sciences | 4 | 2 (50%) |
| Psychological Medicine | 17 | 13 (76%) |
| Clinical Psychological Science | 15 | 9 (60%) |
| Bipolar Disorders | 15 | 6 (40%) |
| Acta Psychiatrica Scandinavica | 11 | 7 (64%) |

Values are n, or n (% of all articles). Screening was performed to exclude articles that were not full-length original research articles (e.g. reviews, editorials, perspectives, commentaries, letters to the editor, short communications, etc.), were not published in March 2020, or did not have a methods section. No issue indicates that the journal did not publish an issue or any articles in March 2020.
\* Journals were included on both the neuroscience (Table S1) and psychiatry lists.

**Table S4:**
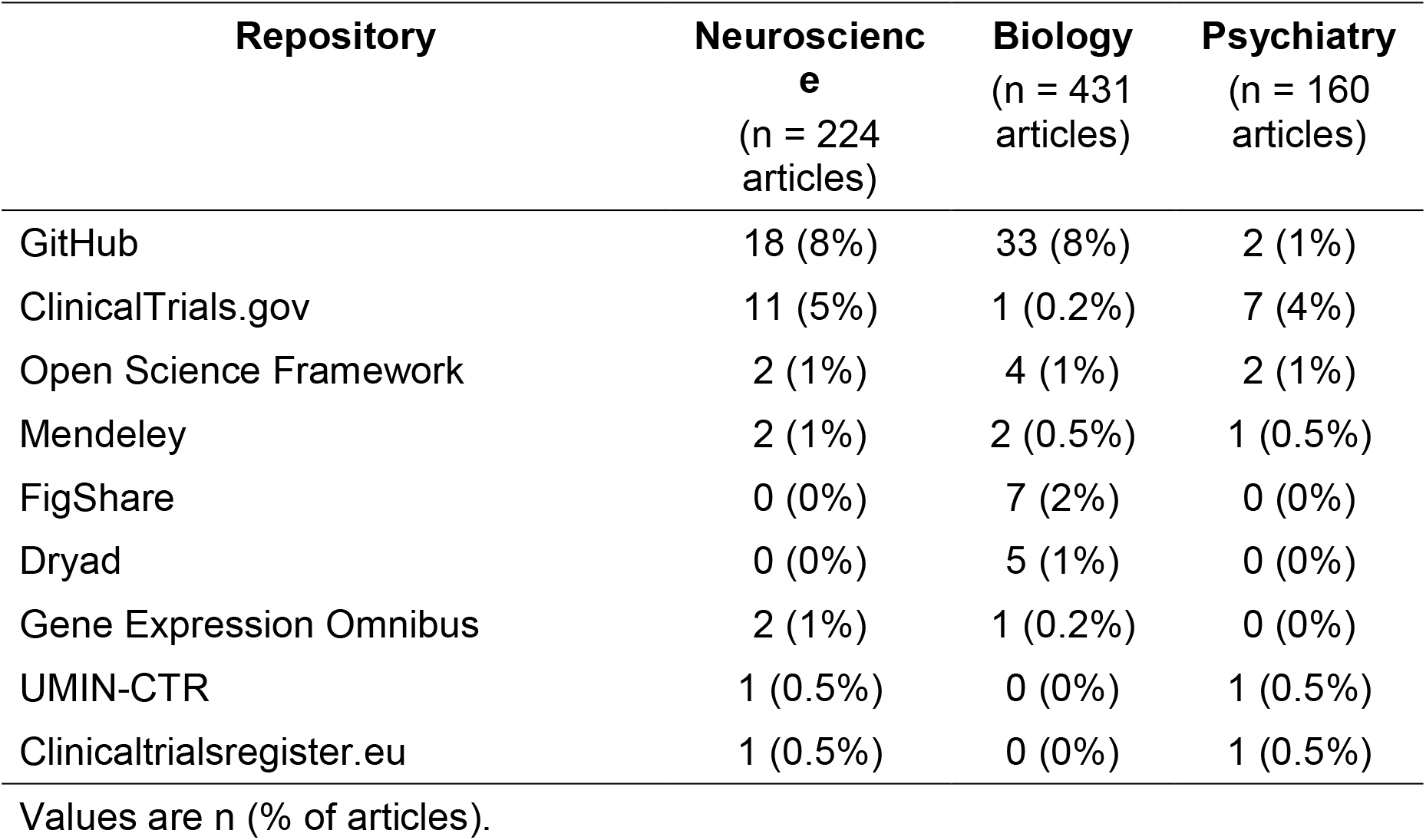
Methods repositories used in neuroscience, biology and psychiatry Values are n (% of articles).

| **Repository** | **Neuroscience**<br>(n = 224 articles) | **Biology**<br>(n = 431 articles) | **Psychiatry**<br>(n = 160 articles) |
| --- | --- | --- | --- |
| GitHub | 18 (8%) | 33 (8%) | 2 (1%) |
| ClinicalTrials.gov | 11 (5%) | 1 (0.2%) | 7 (4%) |
| Open Science Framework | 2 (1%) | 4 (1%) | 2 (1%) |
| Mendeley | 2 (1%) | 2 (0.5%) | 1 (0.5%) |
| FigShare | 0 (0%) | 7 (2%) | 0 (0%) |
| Dryad | 0 (0%) | 5 (1%) | 0 (0%) |
| Gene Expression Omnibus | 2 (1%) | 1 (0.2%) | 0 (0%) |
| UMIN-CTR | 1 (0.5%) | 0 (0%) | 1 (0.5%) |
| Clinicaltrialsregister.eu | 1 (0.5%) | 0 (0%) | 1 (0.5%) |

Values are n (% of articles).

**Table S5:** Number of probable and possible shortcut citations within a paper for each field.

| Field | Number of probable<br>shortcut citations per<br>paper | Number of possible shortcut<br>citations per paper |
| --- | --- | --- |
|  | Median (IQR) | Median (IQR) |
| Neuroscience | 3 (1, 6) | 2 (1, 4) |
| Biology | 2 (1, 4) | 1 (0, 2) |
| Psychiatry | 3 (1, 5) | 2 (1, 4) |

This table shows summary statistics for Figure 2b. Abbreviations: IQR, interquartile range.

**Table S6:** Age of the youngest and oldest shortcut citations within a paper for each field.

| **Citation age** | **Field** | **Probable shortcut citations** | **Possible shortcut citations** |
| --- | --- | --- | --- |
|  |  | Median (IQR) in years | Median (IQR) in years |
| Youngest | Neuroscience | 4 (2, 6) | 3 (2, 6) |
|  | Biology | 3 (2, 6) | 4 (2, 7) |
|  | Psychiatry | 4 (2.75, 7.25) | 4 (2, 7) |
| Oldest | Neuroscience | 21 (10.5, 33) | 14 (7, 22) |
|  | Biology | 14 (7, 23) | 10 (5, 16) |
|  | Psychiatry | 24.5 (11, 35.25) | 19 (10, 30) |

This table shows summary statistics for Figure 3a. Abbreviations: IQR, interquartile range.

**Table S7:** Comparing methods of sharing detailed protocols.

| Characteristic | Shortcut citation | Supplemental files of paper | Protocol repository | Protocol journal |
| --- | --- | --- | --- | --- |
| Examples | N/A | N/A | protocols.io<br>Protocol Exchange<br>ClinicalTrials.gov (clinical trials)<br>PROSPERO (systematic review pre-registration) | **Online:** Bio-protocol.org<br>**Print:** Nature Protocols, “Current Protocols in ...” series<br>**Video:** JOVE |
| Structured format | N/A | No<br>No consistency in what is reported; typically follows the overview format of methods sections; step-by-step protocols are rare | Yes<br>Platforms differ in the level of detail required (essential information vs. comprehensive, step-by-step protocols) | Yes<br>Step-by-step protocols<br>Video protocols |
| Tracks protocol evolution (Are updates possible?) | Citing author must specify modifications to cited methods | **Static:** Can’t be updated after publication | **Living protocol:** Authors can quickly share updated versions. Repositories such as protocols.io allow authors to share forked versions specifying their modifications of someone else’s protocol. Versioning and forking allow scientists to track changes over time, both within and across labs. | **Static:** Reflects what one lab is doing at a single point in time |
| Peer reviewed | Cited resource may or may not be peer reviewed | Supplements may or may not be peer-reviewed, depending on the journal | No, but versions or forks indicate re-use<br>* | Yes |
| Effort required | Low | Low to medium | Medium | High |
| DOI citable | N/A | Yes (DOI for paper) | Yes | Yes |
| Findable | N/A | No<br>Search engines can only find the paper, which may or may not include supplements | Yes<br>Scientists can search protocol repositories | Yes<br>Indexed in major publication search engines |
| Open access (OA) | Cited resource may or may not be OA | Depends on the journal / publisher. Some paywalled publishers offer free access to supplements; others don't. Supplemental files aren't consistently shared on publication repositories or through interlibrary loan programs. | OA | Depends on the journal and article |
| Other considerations | Publications with detailed methods descriptions that are similar to the authors' protocols may not be available |  | Protocols remain available and findable over time, even if the research group moves or the scientist responsible for the protocol has left.<br>Some repositories can be used to share protocols at any time, whereas others are designed to pre-register protocols before the study begins. | Often used to share and obtain credit for new, innovative methods<br>Some clinical journals publish clinical study protocols for ongoing clinical studies |

Abbreviations: JOVE, Journal of Visualized Experiments; N/A, not applicable; OA, open access
\*protocols.io offers a partnership with PLOS One where authors can publish a methods article linked to their protocol

## Notes

https://osf.io/d2sa3/

### Competing Interest Statement

The authors have declared no competing interest.

